# Identification of chemotherapy targets reveals a nucleus-to-mitochondria ROS sensing pathway

**DOI:** 10.1101/2023.03.11.532189

**Authors:** Junbing Zhang, Claire M. Simpson, Jacqueline Berner, Harrison B. Chong, Jiafeng Fang, Zehra Ordulu Sahin, Tom Weiss-Sadan, Anthony P. Possemato, Stefan Harry, Mariko Takahashi, Tzu-yi Yang, Marianne Richter, Himani Patel, Abby E. Smith, Alexander D. Carlin, Adriaan F. Hubertus de Groot, Konstantin Wolf, Lei Shi, Ting-Yu Wei, Benedikt R. Dürr, Nicholas J. Chen, Tristan Vornbäumen, Nina O. Wichmann, Venkatesh Pooladanda, Yuske Matoba, Shaan Kumar, Eugene Kim, Sara Bouberhan, Esther Olivia, Bo Rueda, Nabeel Bardeesy, Brian Liau, Michael Lawrence, Matt P. Stokes, Sean A. Beausoleil, Liron Bar-Peled

## Abstract

Multiple chemotherapies are proposed to cause cell death in part by increasing the steady-state levels of cellular reactive oxygen species (ROS). However, for most of these drugs exactly how the resultant ROS function and are sensed is poorly understood. In particular, it’s unclear which proteins the ROS modify and their roles in chemotherapy sensitivity/resistance. To answer these questions, we examined 11 chemotherapies with an integrated proteogenomic approach identifying many unique targets for these drugs but also shared ones including ribosomal components, suggesting one mechanism by which chemotherapies regulate translation. We focus on CHK1 which we find is a nuclear H_2_O_2_ sensor that promotes an anti-ROS cellular program. CHK1 acts by phosphorylating the mitochondrial-DNA binding protein SSBP1, preventing its mitochondrial localization, which in turn decreases nuclear H_2_O_2_. Our results reveal a druggable nucleus-to-mitochondria ROS sensing pathway required to resolve nuclear H_2_O_2_ accumulation, which mediates resistance to platinum-based chemotherapies in ovarian cancers.

## Introduction

ROS represent a distinct family of reactive molecules that arise during normal cellular metabolism and are further generated in the context of disease states or toxin exposure (Chio and Tuveson, 2017). The reactive nature of these molecules allows them to exert substantial control over multiple cellular pathways through the direct modification of proteins, nucleic acids, or lipids (Chio and Tuveson, 2017; Reczek and Chandel, 2015). Accordingly, ROS homeostasis is tightly regulated within cells through compartmentalization and detoxification. With regards to compartmentalization, mitochondria are best appreciated for their functions as intracellular ROS generators and sinks (Chouchani et al., 2017; Collins et al., 2012). Importantly, ROS levels are thought to dictate the scope of their cellular targets. Low levels of ROS are required to maintain normal cellular homeostasis, functioning in numerous signaling capacities through the modification of phosphatases and metabolic enzymes. At high levels, ROS damage nucleic acids, inactivate proteins and induce lipid peroxidation, leading to ferroptotic cell death (Dixon and Stockwell, 2019). In this regard, chemotherapies that increase the steady-state levels of ROS are currently used in the treatment of multiple cancers, including lymphoma (arsenic trioxide, ATO), ovarian (cisplatin, CPT), bladder (doxorubicin, DOXO) and pancreatic cancers (5-fluorouracil, 5FU)(Chio and Tuveson, 2017; Perillo et al., 2020; Yang et al., 2018). However, the mechanisms underlying ROS increase following chemotherapy, their sensing and targets are poorly described (Chabner and Longo, 2011). As a result of this knowledge gap, there is limited understanding of chemoresistance, one of the greatest clinical challenges in modern cancer treatment and the promise of harnessing ROS as therapeutic modality remains to be fully realized.

Whereas the non-specific nature of heightened ROS levels following chemotherapy treatment and its destruction of DNA is proposed to underlie much of the activity of these drugs (Chabner and Longo, 2011), there is a growing appreciation that ROS modification of specific proteins involved in key cellular pathways may also contribute to chemotherapy cytotoxicity (Liou and Storz, 2010). Elucidating the mechanism of action of these ROS and in particular their molecular targets, has historically been problematic given the transient nature of ROS, the high concentrations required to observe phenotypic changes and the use of non-specific readouts, which have greatly limited functional insights. The global dissection of ROS-target proteins has been advanced using chemical proteomic technologies to profile changes in the reactivity of the amino acid, cysteine (Backus, 2019). The unique chemical property of cysteine makes this residue a primary ROS target with central roles in regulating protein function. Using electrophilic cysteine-reactive probes, recent studies have categorized the direct cysteine targets of H_2_O_2_ and electrophilic lipids in vitro, revealing that ROS-regulated cysteines exist in kinases and metabolic enzymes (Wang et al., 2014). Moreover, these platforms have been used to define the mechanisms by which cells reprogram their redox environment to protect essential pathways (Bar-Peled et al., 2017; Chio et al., 2016; Xiao et al., 2020).

While these chemical proteomic studies provide a list of ROS targets, a key challenge is to functionally understand how these protein targets contribute to the phenotypic consequences of ROS following chemotherapy treatment. To address these challenges, we describe an integrated approach comprised of cysteine-based chemical proteomics and functional genomic CRISPR screening. Using this framework, we provide a comprehensive portrait of functional protein targets of ROS following treatment with 11 different chemotherapies. We identified distinct proteins targeted by each cytotoxic agent, with principal targets including ribosomal proteins, which we connect as regulatory sites for chemotherapy regulated translation. By leveraging this integrated approach, we uncover a nucleus-based ROS sensor that controls compartmentalized H_2_O_2_ levels through regulation of mitochondrial translation. We demonstrate nuclear H_2_O_2_ modifies a functional and conserved cysteine within CHK1. This modification leads to CHK1 activation throught a conformation change in its autoinhibitory domain, leading to the activation of the kinase which launches a cellular program required to decrease nuclear H_2_O_2_ levels. Functional studies using a clinical CHK1 inhibitor, delineated a nuclear-to-mitochondria ROS sensing pathway that couples DNA damage response to control of mitochondrial translation, through the regulation of the mitochondrial DNA (mtDNA) binding protein SSBP1. Loss of SSBP1 decreases nuclear H_2_O_2_ levels and provides resistance to platinum-based chemotherapies in ovarian cancer models. Our findings underscore the value of integrating distinct reads outs of ROS activity to systematically characterize the cellular response to chemotherapies and precision-oncology medicines.

## Results

### Chemotherapies regulate cysteine reactivity

We sought to investigate functionally relevant protein targets of ROS following chemotherapy treatment. Our studies focused on multiple chemotherapies at different stages of clinical evaluation that have been shown to increase steady-state ROS levels (**Supplementary Table 1**). These agents potently blocked the growth of the K562 cell line at 4 days using previously a clinically relevant dose range as previously reported for these anti-cancer compounds (**Figure S1A, Supplementary Table 1**). Testing these agents at ∼5X their IC_50_ concentrations revealed minimal loss of cell viability at a 24 hr. time period, providing a context for exploring ROS signaling independent of secondary effects resulting from proliferation arrest (**Figure S1B**). We characterized changes in the redox status of K562 cells following chemotherapy by measuring the intensity of 2’,7’-dicholorodihydrofluourscien diacetate (DCF) and the levels of metabolites and pathways required for ROS detoxification including: NAD+/NADH (Wu et al., 2016) NADP+/NADPH (Wang et al., 2017) and NRF2 signaling (**Figures 1A**, **S1C**). This analysis revealed both time and chemotherapy-specific differences in redox response pathways following drug treatment.

**Figure 1:**
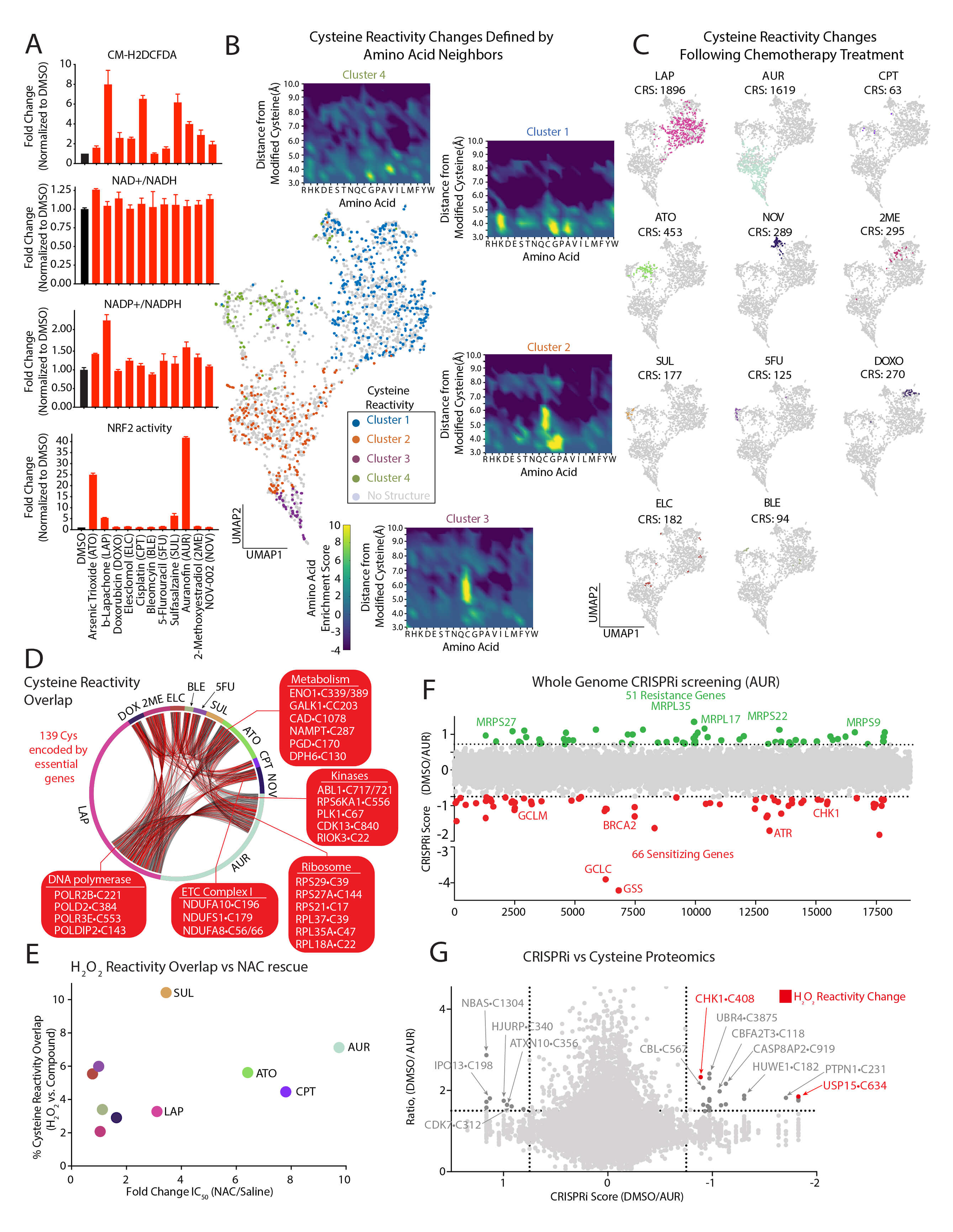
Proteogenomic characterization of chemotherapy targets. (A) Chemotherapies regulate steady state levels of ROS and antioxidant response pathways. Changes in DCF intensity, NAD+/NADH, NADP+/NADPH, and NRF2 activity in K562 cells following treatment with 5.1µM ATO, 1.5µM LAP, 0.1µM DOXO, 30nM ELC, 8.3µM CPT, 7.1µM BLE, 6.0µM 5FU, 0.5mM SUL, 2.5µM AUR, 1µM 2ME or 0.5mM NOV for 24 hrs (see also **Figure S1B).** (B) Chemotherapy-mediated cysteine reactivity changes are marked by distinct structural features. Commonly detected cysteines regulated by chemotherapies are found in four distinct clusters and are color-coded based on available structures as depicted in the UMAP (center). Enrichments plots of residues within a 10Å radius of reactive cysteines were determined for each cluster (see methods). Cysteine reactivity changes were determined by isoTMT and were included as part of the UMAP if their ratio (R) of reactivity change following treatment was R>1.5 and was identified in all proteomic runs (see methods, **Supplemental Tables 2,9**). (C) UMAP of cysteine reactivity changes following chemotherapy treatment. Cysteine reactivity score (CRS) was determined as described in the methods (see also **Figure S2D**). (D) Connectivity diagram for shared cysteine reactivity changes following chemotherapy treatment. (E) Comparison of each chemotherapy’s overlap with H_2_O_2_ cysteine reactivity and fold-change in IC_50_ following NAC treatment (see also **Figure S1C, Supplemental Table 7**). (F) Genome-wide CRISPRi screen in K562 cells identifies genes that mediate sensitivity and resistance to AUR. Positive CRISPR score (in green) represent genes whose loss provides resistance to AUR and negative CRISPR scores (in red) represent genes whose loss sensitizes cells to AUR (see also **Supplemental Table 5**). (G) Comparison of targets from AUR CRISPRi screen with AUR iso-TMT screen. Genes whose corresponding cysteines have a decrease in reactivity following H_2_O_2_ treatment are marked in red (see also **Supplemental Tables 5,7**).

We next sought to identify proteins targeted by chemotherapies by monitoring changes in cysteine reactivity using the iso-TMT platform (Bar-Peled et al., 2017; Kuljanin et al., 2021; Vinogradova et al., 2020). We rationalized that using this unbiased approach would allow us to capture the largest number of cysteine modifications following chemotherapy treatment, including oxidation, direct modification by a compound and adduction by ROS-related metabolites (e.g., lipid peroxides). These modifications are collectively read out as changes in cysteine reactivity. We analyzed cells treated at early time points (2-3 hrs.) to minimize changes in cysteine reactivities brought about by expression changes. We defined chemotherapy-based cysteine reactivity changes as those showing an iso-TMT Ratio (R) with ≥1.5-fold change in reactivity compared to vehicle control. Out of 35656 cysteines and 8297 proteins identified, we found that 4980 cysteines within 2910 proteins had changes in their reactivity (**Supplementary Table 2**). K-means clustering of a subset (2498) of reactive cysteines detected in all proteomic experiments resulted in four distinct clusters of reactive cysteines (**Figure 1B**). To characterize the structural features underlying this clustering, we analyzed protein structures containing 4000+ commonly detected reactive and non-reactive cysteines (**Supplemental Table 9**). By identifying the nearest amino acid neighbors within a 10Å sphere centered on each cysteine of interest, we found specific amino acids enriched in each cluster (**Figures 1B, S2A-B**). For example, proximal cysteines are strongly selected for in clusters 2 and 3, suggesting the presence of disulfide bonds upon oxidation of the corresponding reactive cysteine. Interestingly, we identified a proximal lysine in Cluster 1, suggesting the presence of the recently described lysine–cysteine redox switch (Wensien et al., 2021) (**Figures 1B, S2B**). We found that amino acids identified by our structural analysis were not encapsulated in the primary sequence surrounding a cysteine of interest, implying that the chemical properties of cysteine reactivity changes may be missed by analyzing primary sequences alone (**Figure S2C**). Using commonly detected cysteines, we developed a ‘cysteine reactivity score’, finding that AUR- and LAP-reactive cysteines had the greatest changes in cysteine reactivity among all treatments (**Figures 1C, S2D**). By concentrating our analysis on cysteines identified in all treatments, we found 500+ common cysteine targets regulated by two or more agents (**Figure 1D**) in addition to cysteines that were distinctly targeted by each drug (**Figure 1C**). The cysteine reactivity score correlated with DCF staining, with the notable exception of SUL (**Figure S2E**). This suggests that the cysteine reactivity score may provide a faithful representation of the cellular redox status.

We found that modified cysteines mapped to multiple pathways, including protein translation, glycolysis, DNA replication, and mTORC1 nutrient sensing (Figure **S2F, Supplementary Table 3)**. Multiple ribosomal cysteines contained chemotherapy-sensitive cysteines (**Figures S2F, S3A**), suggesting a mechanistic link between changes in cysteine reactivity and translational control. Treatment of cells with chemotherapies that modified ribosomal proteins revealed that AUR, CPT, 2ME and to a lesser extent, decreased protein synthesis, whereas chemotherapies that do not modify ribosomal cysteines (e.g., BLE, 5FU and SUL) did not have a measurable effect (**Figure S3B**). Among the chemotherapy regulated ribosomal cysteines, we find that C22 in RPL18A and C39 in RPLA37A were altered following treatment with multiple agents, implicating these proteins as potential ribosomal sensors of redox imbalance (**Figure S3C**). We determined that 82% of cysteine reactivity changes reflected bona-fide changes in reactivity, while we attribute the remaining changes to alterations in protein expression (**Supplemental Table 2**). For AUR treatment, we monitored reactivity changes at 2 and 6 hrs. to test if any early changes in cysteine reactivity would translate to changes in protein abundance. To this end, we overlaid modified cysteines with known ubiquitination sites on proteins and discovered that 41% of proteins whose expression was reduced contain a ubiquitination site within ≤20 residues of a modified cysteine (**Figure S2G**). For example, in AKT2, a critical regulator of cell growth and metabolism, C297 is the only cysteine modified at early time points. However, at later time points multiple cysteines on AKT2 are modified, decreasing expression by 1.5-fold, an observation that we confirmed by immunoblot (**Figure S2H**)– suggesting the existence of degrons specific to ROS-based cysteine modification. Collectively, these findings provide a comprehensive portrait of chemotherapy regulated cysteines and the immediate cellular pathways that are impacted and may respond to this ROS.

### Defining mechanisms of sensitivity and resistance to ROS regulated by chemotherapy treatment

Because cysteine reactivity changes can encompass many different modifications, we sought to prioritize agents that operate via an explicit ROS-based mechanism. For each compound, we compared its cysteine reactivity changes to those mediated by H_2_O_2_ (**Supplemental Table 7**) to the change in cytotoxicity following treatment with n-acetyl cysteine (NAC), a molecule with ROS scavenging abilities (Zhitkovich, 2019). This comparison highlighted AUR as having the greatest overlap with H_2_O_2_ targets and rescue by NAC (**Figure 1E**), suggesting that it could be relevant to cellular pathways that sense and respond to an increase in H_2_O_2_. AUR is an FDA-approved drug for rheumatoid arthritis and is currently under experimental investigation for the treatment of chronic lymphocytic leukemia (NCT01419691) and ovarian cancer in combination with Sirolimus (NCT03456700, 2018). Although the established targets of this gold thiolate are TXNRD1/2 (Stafford et al., 2018), enzymes which are critical in the cellular antioxidant response, we suspected that additional pathways may contribute to the cellular response to AUR, given the diversity of its cysteines targets. Importantly, we believe that AUR can be used as a tool to help understand the ROS-based pathways which are targeted by other chemotherapies.

To this end, we performed a genome-wide CRISPRi screen to identify mechanisms of sensitivity and resistance to AUR treatment. K562 cells expressing dCas9-KRAB (Gilbert et al., 2014) were infected with a genome-wide sgRNA library and grown for 11 population doublings in the presence of vehicle or 1 µM of AUR (**Figure 1F, Supplemental Table 5**). For each gene, we calculated a CRISPRi score by comparing the relative fold change between corresponding sgRNAs enriched in AUR vs. vehicle. This analysis identified 51 genes mediating resistance (e.g., whose corresponding sgRNAs were enriched in the treated cells) and 66 genes mediating sensitivity (e.g., sgRNAs depleted in the treated cells). The subcellular localization of hits from this screen highlighted the mitochondria and the nucleus as hubs of resistance and sensitivity, respectively (**Figure S3E**). We found multiple cellular pathways correlating with resistance, including those required for mitochondrial translation and ETC complex assembly. In contrast, disruption of the DNA damage response (DDR) and the glutathione biosynthetic pathway (Harris et al., 2019) strongly sensitized cells to AUR treatment, with the latter confirming the robustness of this study (**Figures S3D,F)**.

To prioritize AUR targets for mechanistic characterization, we calculated a ‘prioritization score’ by integrating the CRISPRi score and cysteine reactivity change for a given target (**Figure 1G**, **Supplemental Table 6**). We reasoned that relevant targets would belong to cellular networks whose members are likely to mediate AUR cytotoxicity or become modified following treatment. Thus, we incorporated the 10 closest interactors as defined by STRING (Jensen et al., 2009) into our prioritization score. This approach revealed multiple target proteins and corresponding networks, with an enrichment for high priority AUR targets/networks belonging to stress response or E3-ubiqutin ligase pathways (**Figures S4A-B**). Many of these pathways have been previously connected to ROS control (Arnandis et al., 2018; Gajewska et al., 2021; Niederkorn et al., 2022; Salmeen et al., 2003; Zarei et al., 2019) and localize to the nucleus (**Supplemental Table 4**). To further focus our analysis on response to ROS, we overlaid H_2_O_2_-mediated changes in cysteine reactivity within the AUR response networks, revealing that the DNA damage kinase CHK1 (Ciccia and Elledge, 2010) is required for mediating sensitivity to AUR and is further modified by H_2_O_2_ (**Figures 1G, S4C-E**). This finding suggests that CHK1 may participate in nuclear ROS sensing and response.

### Identification of CHK1•C408 as a nuclear ROS sensor

CHK1 is modified at C408, a highly conserved cysteine that lies within the C terminal KA1 domain (**Figure 2A**). KA1 functions in an autoinhibitory capacity by binding and blocking the activity of the N terminal CHK1 kinase domain (Katsuragi and Sagata, 2004). Short-term treatment with AUR led to CHK1 activation as measured by CHK1 autophosphorylation at S296, prior to a change in the phosphorylation of H2AX•S139 (**Figure 2B**), an established marker of DNA damage. Importantly, AUR activation of CHK1 was blocked by treatment with the antioxidant NAC (**Figure 2B**). To test the hypothesis that nuclear H_2_O_2_ may be involved in CHK1 activation, we first confirmed that AUR treatment increased nuclear H_2_O_2_ levels using two different nuclear localized H_2_O_2_ reporters, HyPer7 and roGFP2-Orp1 (Gutscher et al., 2009; Pak et al., 2020) (**Figure S5A**). Both reporters were oxidized following treatment with AUR, and this oxidation was completely rescued following treatment with NAC (**Figures 2C, S5B-C**). Treatment of cells with H_2_O_2_ led to an increase in binding of the sulfinic acid specific probe, nitroso-desthiobiotin (NO-DTB) (Lo Conte et al., 2015; Reddie et al., 2008; Yang et al., 2016), to CHK1•C408 (**Figure 2D**). There was a concomitant decrease in the labeling of CHK1•C408 by IA-DTB, suggesting that H_2_O_2_ directly oxidizes CHK1 (**Figures 2D, S5D**). To determine if CHK1•C408 oxidation by H_2_O_2_ leads to kinase activation, we targeted D-amino acid oxidase (DAAO) to the nucleus (**Figure S5E**). DAAO is an enzyme which oxidizes D-amino acids to their corresponding *α*-keto acids producing H_2_O_2_ (Steinhorn et al., 2018) and short-term treatment with D-Ala but not L-Ala leads to an increase in CHK1 activity in a NAC-dependent manner (**Figure 2E**). At early time points, activation of nuclear DAAO did not increase H2AX•S139 phosphorylation (**Figure 2E**), suggesting that H_2_O_2_ activation of CHK1 precedes activation of canonical DNA damage markers. By fusing DAAO to the N-terminus of CHK1 and localizing the chimeric protein to the mitochondrial outer membrane, we found H_2_O_2_ is sufficient to activate CHK1 in cells (**Figures 2F-G**). Finally, in vitro treatment of CHK1 with H_2_O_2_ led to a dose-dependent increase in CHK1 activity (**Figures 2H, S5F**). Using a CHK1 in vitro binding assay, we found that that addition of H_2_O_2_ strongly diminished the interaction between the KA1 and CHK1 kinase domain (**Figure 2I**), providing a mechanism by which CHK1•C408 oxidation activates CHK1.

**Figure 2:**
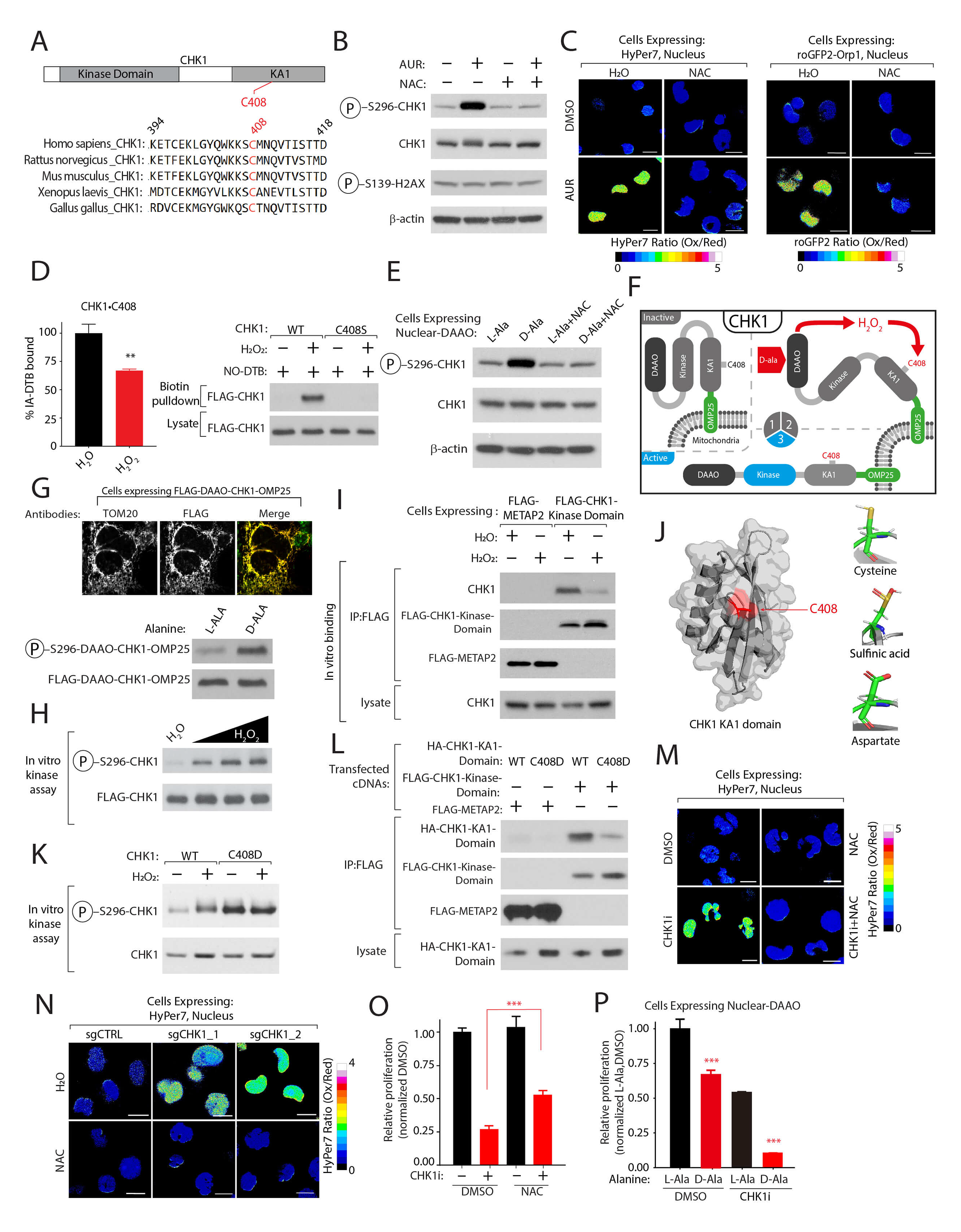
CHK1 functions as a sensor and regulator of nuclear H_2_O_2_ levels. (A) C408 is highly conserved. ClustalW alignment of CHK1 KA1 domain from the indicated species. (B) AUR activates CHK1 in a NAC dependent manner. K562 cells were treated with AUR in the presence of NAC or vehicle control. CHK1 activity was determined by immunoblotting for the indicated proteins. (C) AUR increases the steady-state levels of nuclear H_2_O_2_. Ratiometric images (Oxidized(Ox)/Reduced(Red)) of HyPer7 or roGFP2-ORP1 H_2_O_2_ reporters localized to the nucleus following treatment with DMSO or 1.5 µM AUR in K562 cells in the presence or absence of 5mM NAC (see also **Figures S5A-C**). (D) H_2_O_2_ treatment regulates CHK1•C408 oxidation. Left, iso-TMT ratio of CHK1•C408 following treatment of K562 cells with 200 µM H_2_O_2_ (see also **Supplemental Table 7**). Right, HEK-293T cells expressing FLAG-CHK1 or CHK1•S408S and treated with 100 µM H_2_O_2_,. Immunoblot analysis of sulifinic acid modification on CHK1 following enrichment from lysates treated with a biotin-containing sulfinic acid probe, NO-DTB (see methods). (E) Nuclear H_2_O_2_ activates CHK1. K562 cells stably expressing D-amino acid oxidase (DAAO) localized to the nucleus were treated for 3 hrs. with either 10mM L-Ala or D-Ala or 5mM NAC and CHK1 kinase activity was determined by immunoblot as described in (B). (F) Schematic depicting the localization of DAAO-CHK1 to the mitochondrial outer membrane. (G) H_2_O_2_ is sufficient to activate CHK1. Top, Immunofluorescence image of DAAO-CHK1-OMP25 in HEK- 293T cells co-stained with mitochondrial resident protein TOM-20. Bottom, DAAO-CHK1-OMP25 kinase activity was determined by immunoblot following treatment with either 10mM L-Ala or D-Ala as described in (B). (H) H_2_O_2_ directly activates CHK1. Representative analysis of CHK1 in vitro activity following treatment with increasing amounts of H_2_O_2_. (I) H_2_O_2_ reduces the interaction between endogenous CHK1 and the CHK1 kinase domain. Lysates from K562 cells expressing FLAG-tagged METAP2 (control) or CHK1 kinase domain were treated H_2_O or 100 µM H_2_O_2_ Following immunoprecipitation with anti-FLAG M2 beads, interaction with endogenous CHK1 was determined by immunoblot. (J) Right, Crystal structure of C-terminal Kinase Associate 1 (KA1) domain of CHK1 highlighting the location of C408 in red, adapted from PDB ID: 5WI2 (Chen et al., 2000). Left, Modeling of CHK1 C408 interactions as a sulfinic acid or when mutated to Asp. (K) CHK1•C408D has high levels of kinase activity in vitro and cannot be activated further following addition of 1000 µM H_2_O_2_. (L) CHK1•C408D-mutation in KA1 domain blocks interaction with CHK1 kinase domain. Immunoblot analysis of the interaction between KA1-CHK1 kinase domain was determined following immunoprecipitation of FLAG-tagged proteins from lysates generated from HEK-293T cells expressing the indicated proteins. (M-N) CHK1 inhibition increases steady-state nuclear H_2_O_2_ levels. Ratiometric (Ox/Red) images of HyPer7 localized to the nucleus in K562 cells following 48 hrs treatment with 2 µM CHK1 inhibitor MK-8776 (CHK1i) (M) or in K562-dCas9-KRAB cells expressing sgCTRL or sgCHK1 (N). (O) NAC treatment partially rescues CHK1i cytotoxicity. K562 cells were co-treated with 1 µM CHK1i and 5mM NAC and relative proliferation was determined after 96 hrs. by measuring cellular ATP concentrations. (P) Nuclear H_2_O_2_ increases CHK1i cytotoxicity to block cell proliferation. K562 cells stably expressing nuclear localized DAAO were pretreated for 72 hrs. with 10mM L-Ala or D-Ala prior to treatment with vehicle or 0.5 µM CHK1i. Relative proliferation was determined as described in (O). Scale Bar=10 µm. **p < 0.001, ***p< 0.0001.

Although no amino acid mutation could mimic C408 oxidation, we mutated this residue to Asp to model an oxidized form of cysteine, sulfinic acid (Permyakov et al., 2012) (**Figure 2J**). In vitro, the CHK1•C408D mutant and H_2_O_2_-treated CHK1 demonstrated equivalent activation. However, the mutant could not be further activated, indicating that C408 is the target of H_2_O_2_ in the kinase (**Figure 2K**). Using a modified binding assay that monitors KA1 and kinase domain interaction, we confirmed that the KA1 domain harboring the C408D mutant poorly interacted with the N-terminal kinase domain of CHK1 (**Figure 2L**). Following AUR treatment, we found that K562 cells stably expressing FLAG-CHK1•C408D proliferate to a greater degree than cells expressing a control protein (METAP2) or WT CHK1, with a concomitant reduction in H2AX•S139 phosphorylation (**Figures S5G-H**). These data strongly suggest that C408 in CHK1 is a nuclear H_2_O_2_ sensor, which functions by disrupting the association of KA1 domain with the kinase domain leading to CHK1 activation.

### CHK1 inhibition increases steady-state levels of nuclear H_2_O_2_

Given our identification of CHK1 as a nuclear H_2_O_2_ sensor, we wondered whether CHK1 might have a broader role in controlling ROS levels within this compartment. Treatment of K562 cells with a clinical grade CHK1-inhibitor (MK-8776 herein referred to as CHK1i) (Guzi et al., 2011) NCT00779584, 2008) or depletion of CHK1 resulted in a substantial increase in nuclear H_2_O_2_ levels as determined by both the HyPer7 and roGFP2-Orp1 reporters (**Figures 2M-N**, **S6A-E**). Interestingly, treatment with NAC not only rescued CHK1i mediated nuclear H_2_O_2_ but partially rescued proliferation defects and H2AX•S139 phosphorylation incurred by this inhibitor (**Figures 2O**, **S5I**), further supporting the role of CHK1 in ROS response. Consistent with previous reports that demonstrate that high levels of nuclear H_2_O_2_ can inhibit cell growth (Tan et al., 2017), we found that the targeted generation of nuclear H_2_O_2_ with nuclear DAAO enhanced CHK1i-mediated abrogation of cell proliferation (**Figure 2P**). These results suggest that CHK1 is not simply an oxidized bystander, but rather plays an active role in H_2_O_2_ mitigation, with inhibition of this sensor resulting in a corresponding rise in lethal peroxide levels within the nucleus.

### CHK1 phosphorylation of SSBP1 restricts its mitochondrial localization and decreases nuclear H_2_O_2_ levels

Because CHK1 inhibition raises nuclear H_2_O_2_ levels, we hypothesized that a CHK1 substrate might be directly involved in ROS regulation. To identify the substrate, we overlaid CHK1 phosphoproteomics data (Blasius et al., 2011) with our CRISPRi screen to find substrates whose depletion may revert or exacerbate AUR cytotoxicity (**Figure 3A**). Using this approach, we found that SSBP1, a mtDNA binding protein, met the criteria of a potential candidate. SSBP1 is active in the mitochondria, where it facilitates the interaction between mtDNA polymerase pol *γ* and the mtDNA helicase at the mtDNA replication fork (Copeland and Longley, 2003). We verified that reduction in SSBP1 levels led to a pronounced rescue of AUR cytotoxicity (**Figure S6F**). To mechanistically dissect SSBP1 regulation by CHK1, we first established that CHK1 phosphorylates SSBP1 in vitro at S67 (**Figures 3B, S6G**). We found an H_2_O_2_-dependent increase in SSBP1 phosphorylation but did not detect any phosphorylation in a SSBP1•S67A mutant (**Figure 3B**). We also observed the CHK1•C408D mutant had heightened phosphorylation of SSBP1 (**Figure 3C**), consistent with the hyperactive state of this mutant. We confirmed that CHK1 and SSBP1 exist in an epistatic relationship, finding nuclear H_2_O_2_ levels significantly lowered following CHK1i treatment in cells depleted of SSBP1 (**Figures 3D, S6H**).

**Figure 3:**
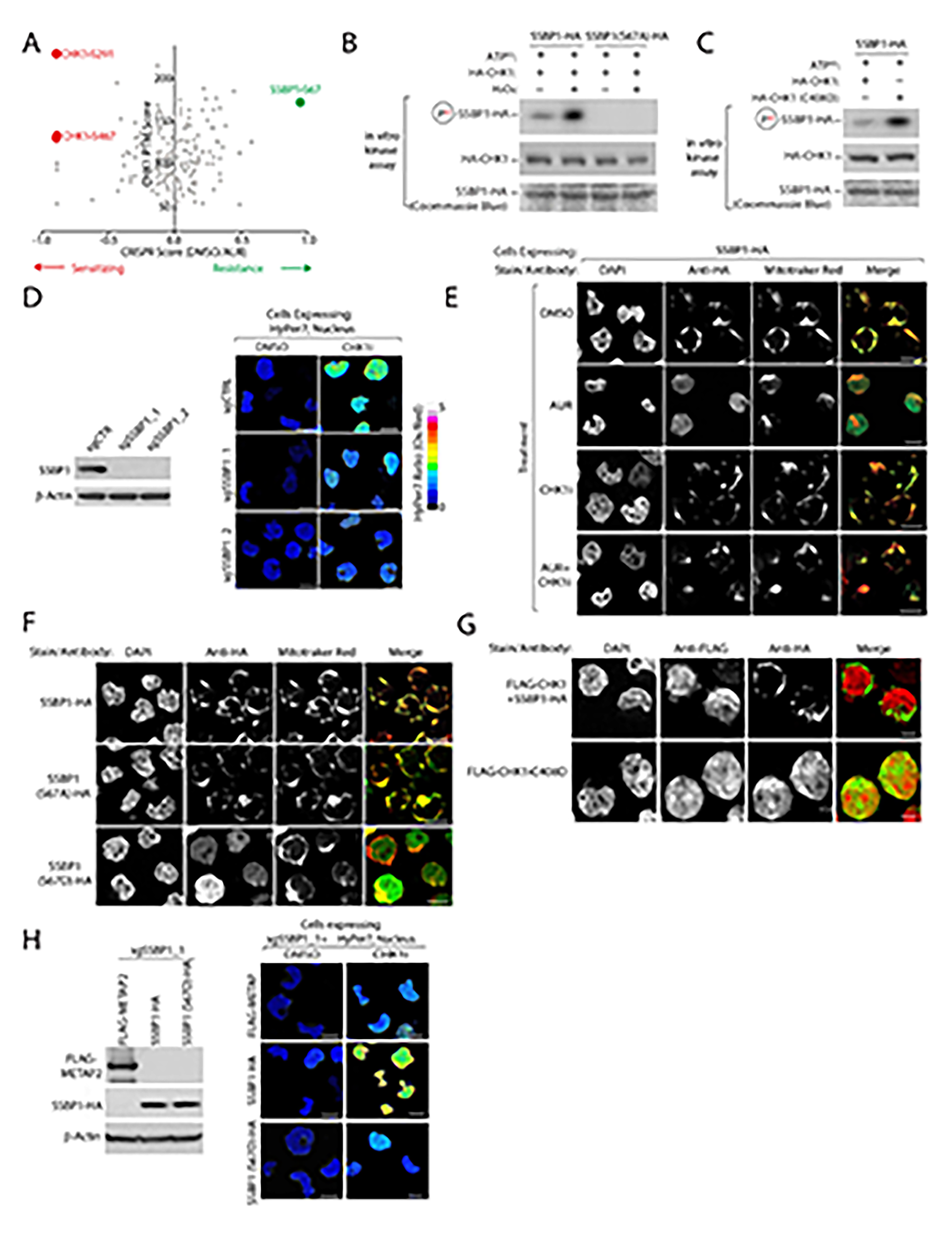
CHK1 phosphorylates SSBP1 blocking its mitochondrial localization to decrease nuclear H_2_O_2_ levels. (A) Comparison of CRISPRi scores following AUR treatment with CHK1 phosphorylation sites characterized in (Blasius et al., 2011) identifies SSBP1•S67 as a potential CHK1 target mediating resistance to AUR. (B) SSBP1 is a direct target of CHK1 that is phosphorylated in a H_2_O_2_-dependent manner. Autoradiogram analysis of CHK1 in vitro kinase assay using purified HA-CHK1 and SSBP1-HA or SSBP1•S67A-HA following treatment with 1000 µM H_2_O_2_. (C) CHK1•C408D is hyperactivated. Autoradiogram analysis of CHK1 in vitro kinase assay as described in (B) using purified HA-CHK1 or HA-CHK1•C408D. (D) SSBP1 regulates nuclear H_2_O_2_ levels downstream of CHK1. Left, immunoblot of SSBP1 levels in K562-dCas9-KRAB cells expressing the indicated sgRNAs. Right, Ratiometric (Ox/Red) images of HyPer7 localized to the nucleus in K562-dCas9-KRAB cells expressing the indicated sgRNAs targeting SSBP1 and treated with vehicle or 2 µM CHK1i. (E) CHK1 phosphorylation blocks SSBP1 mitochondrial localization. The localization of SSBP1-HA was determined by immunofluorescence analysis of K562 cells treated with vehicle, 1.5µM AUR, 2µM CHK1i or AUR/CHK1i for 6 hr. (F) SSBP1•S67D phosphomimetic mutant does not localize to the mitochondria. Immunofluorescence analysis of K562 cells stably expressing the indicated SSBP1-HA proteins. (G) CHK1 is sufficient to drive SSBP1 re-localization. Immunofluorescence analysis of K562 cells stably expressing FLAG-CHK1 or hyperactivated FLAG-CHK1•C408D along with SSBP1-HA. (H) SSBP1•S67D phosphomimetic mutant decreases nuclear H_2_O_2_. Left, reintroduction of SSBP1-HA or SSBP1•S67D-HA into K562 depleted of SSBP1. Levels of the indicated proteins were determined by immunoblot. Right, cells were treated with 2 µM CHK1i or vehicle and nuclear H_2_O_2_ levels were measured with nuclear localized HyPer7. Scale Bar=10 µm.

The mitochondrial localization of SSBP1 is required for its cellular activity (Rajala et al., 2014). Strikingly, we found that CHK1 activation, following AUR treatment, resulted in a diffuse localization of SSBP1 which could be rescued following treatment with CHK1i (**Figure 3E**). A SSBP1•S67D phosphomimetic mutant mirrored the diffuse localization of WT SSBP1 following AUR treatment (**Figure 3F**). The SSBP1•S67A phospho-deficient mutant constitutively localized to the mitochondria even following AUR treatment (**Figure S6I**), suggesting that CHK1 directly regulates SSBP1 localization through CHK1•S67 phosphorylation. Importantly, expression of constitutively active CHK1•C408D resulted in the cytosolic localization of SSBP1, in comparison to cells expressing wildtype CHK1 (**Figure 3G**), indicating that CHK1 activity is sufficient to direct the localization of SSBP1. To evaluate the impact of SSBP1 localization on nuclear H_2_O_2_ levels, we depleted endogenous SSBP1 and co-expressed SSBP1 or SSBP1•S67D, finding that in comparison to WT SSBP1, expression of SSBP1•S67D significantly reduced nuclear H_2_O_2_ levels following CHK1i or AUR treatment (**Figures 3H**, **S6J**). Cells expressing SSBP1•S67D were additionally protected from AUR cytotoxicity relative to cells expressing WT SSBP1(**Figure S6K**), demonstrating that CHK1 regulates nuclear H_2_O_2_ levels through the phosphorylation and subsequent cytosolic retention of SSBP1.

### CHK1-SSBP1 modulates mitochondrial translation to control nuclear H_2_O_2_ levels

Given its role in mitochondrial function, we suspected that SSBP1 may impact mitochondrial ROS, one of the major sites of cellular H_2_O_2_ production (Collins et al., 2012). We found that depletion of SSBP1 led to a significant decrease in mitochondrial matrix H_2_O_2_ and superoxide following treatment with CHK1i (**Figures 4A**, **S7A-B**), suggesting that CHK1 regulates nuclear H_2_O_2_ levels through mitochondrial ROS generation. These results pointed to a nucleus-to-mitochondria signaling pathway, and we found that treatment of cells with Mito-TEMPO, a mitochondrially localized redox modulator (Dikalova et al., 2010), lowered nuclear H_2_O_2_ levels following CHK1i treatment (**Figure 4B**). Moreover, targeted generation of H_2_O_2_ within the mitochondrial matrix was only cytotoxic in the presence of CHK1i (**Figure S7C-D**), suggesting that cytotoxicity imparted by CHK1 inhibition is partially dependent on mitochondrial H_2_O_2_. These epistasis experiments indicate that SSBP1 functions downstream of CHK1 to regulate mitochondrial H_2_O_2_ levels which in turn control nuclear H_2_O_2_ levels.

**Figure 4:**
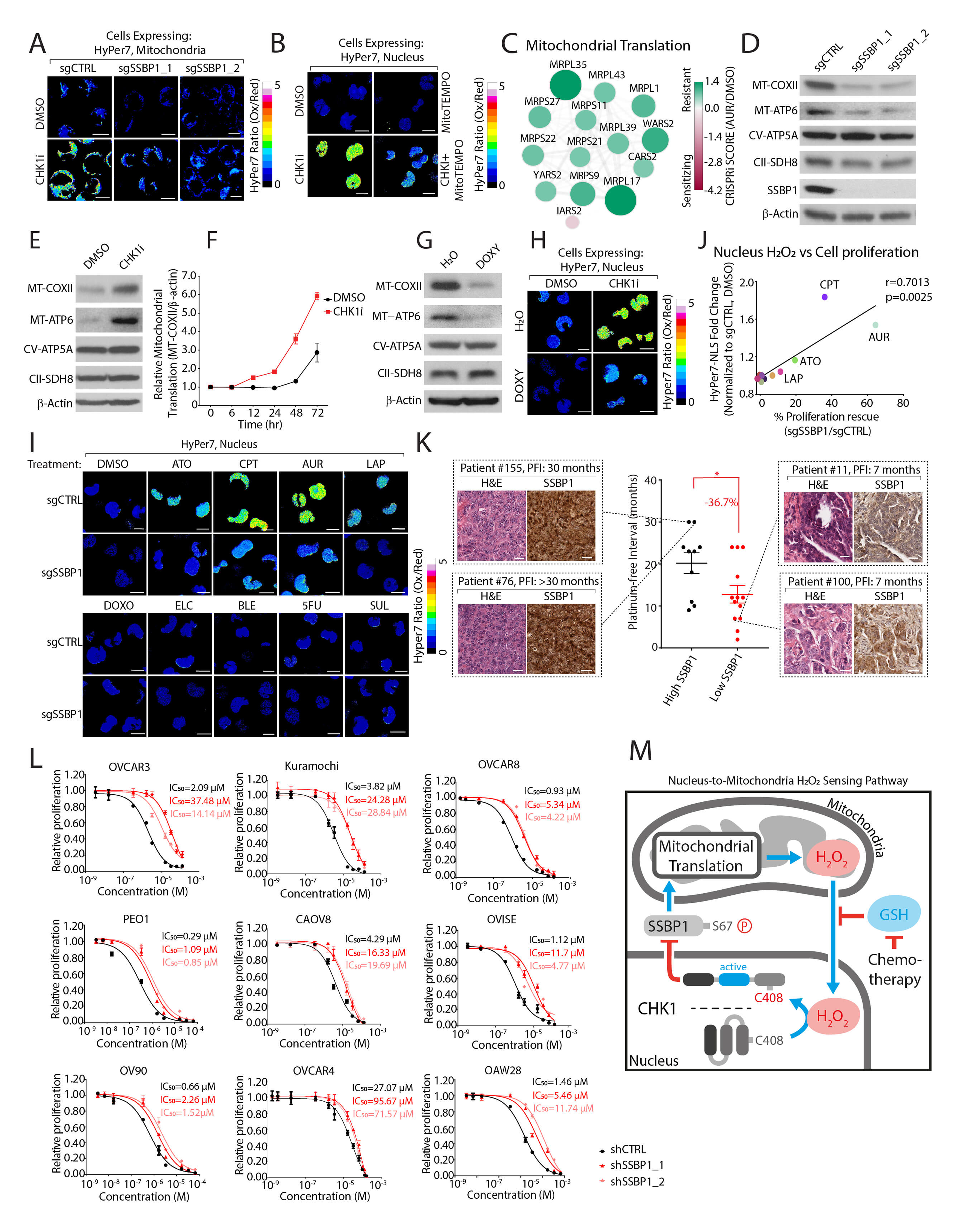
CHK1-SSBP1 regulated mitochondrial translation controls mitochondrial and nuclear H_2_O_2_ levels to mediate cisplatin resistance in ovarian cancer models. (A) CHK1 regulates mitochondrial H_2_O_2_ levels in a SSBP1-dependent manner. Ratiometric (Ox/Red) images of HyPer7 localized to the mitochondrial membrane in K562-dCas9-KRAB cells expressing the indicated sgRNAs and treated with vehicle or 2 µM CHK1i. (B) Mitochondrial H_2_O_2_ levels regulate nuclear H_2_O_2_ levels. Ratiometric (Ox/Red) images of HyPer7 localized to the nucleus in K562 cells treated with 10µM mitoTEMPO and 2 µM CHK1i. (C) Targeting mitochondrial translation mediates AUR resistance. CRISPRi scores of the indicated genes involved in mitochondrial translation. Genes were clustered based on co-essentiality. (D) SSBP1 depletion downregulates mitochondrially translated proteins. Immunoblot analysis of MT-COXII, MT-ATP6 (mitochondrially encoded, top) or ATP5A, UQCRC2 (nuclear encoded, middle) in K562-dCas9-KRAB cells stably expressing the indicated sgRNAs. (E) Inhibition of CHK1 increase mitochondrial translation. K562 cells were treated with 2 µM CHK1i and the levels of mitochondrially translated proteins were determined as in (D). (F) MT-COXII expression following CHK1i treatment. Cells were pre-treated for 48 hrs with 2.3 µM Doxycycline at which time the drug was removed and cells were treated with 2 µM CHK1i or vehicle control. Expression of indicated proteins was determined by immunoblot and normalized to *β*-actin (see also Figure S7F). (G) Doxycycline (DOXY) decreases mitochondrial translation. K562 cells were treated with 2.3 µM DOXY and the levels of mitochondrially translated proteins were determined as in (D). (H) DOXY treatment reduces CHK1i mediated nuclear H_2_O_2_ levels. Ratiometric (Ox/Red) images of K562 cells expressing HyPer7 localized to nucleus and pre-treated for 72 hrs. with 2.3µM DOXY followed by 2 µM CHK1i treatment for 24 hrs. (I) SSBP1 depletion decreases nuclear H_2_O_2_ levels following treatment with chemotherapies. Ratiometric (Ox/Red) images of K562 cells expressing HyPer7 localized to nucleus and treated with the indicated agents. (J) Comparison of fold-change in nuclear H_2_O_2_ levels with proliferation rescue in cells K562 depleted of SSBP1 and treated with the indicated agents. (K) Lower SSBP1 levels correlate with shorter platinum free intervals (PFI) in high-grade serous ovarian cancer (HGSOC) tumors. Insets show representative expression of SSBP1 and H&E staining of HGSOC tumors. (L) Depletion of SSBP1 provides resistance to cisplatin treatment in ovarian cancer cell lines. Cisplatin IC_50_ values were determined in Ovarian cancer cell lines expressing the indicated shRNAs. (M) Model. Nuclear H_2_O_2_ activates CHK1 leading to the phosphorylation and cytosolic retention of SSBP1. Cytosolic SSBP1 cannot promote mitochondrial translation which generates H_2_O_2_. Mitochondrial H_2_O_2_ is transmitted to the nucleus following a decrease in GSH:GSSH ratio by certain chemotherapies.

Recalling that top-scoring resistance genes to AUR toxicity per our CRISPRi screen were involved in mitochondrial translation (**Figures 4C, S7G**), we wondered whether the regulation of nuclear H_2_O_2_ by SSBP1 occurs at the level of mitochondrial translation. Consistent with its regulation of mtDNA (Jiang et al., 2021), depletion of SSBP1 reduced mtDNA (**Figures S7E-F**). and led to a strong downregulation of ETC proteins encoded by mtDNA, including MT-COXII and MT-ATP6 (**Figure 4D**). Other ETC components that are encoded by genomic DNA did not show a similar decrease (**Figure 4D**). Interestingly, inhibition of CHK1 resulted in an increase in MT-COXII and MT-ATP6 expression and further increased mitochondrial translation rates in comparison to vehicle control (**Figures 4E-F**, **S7H**), suggesting that CHK1 indirectly regulates mitochondrial translation through SSBP1. To directly demonstrate that mitochondrial translation is necessary to regulate nuclear H_2_O_2_ levels downstream of CHK1, we treated cells with doxycycline (DOXY), an inhibitor of mitochondrial translation (Chatzispyrou et al., 2015), finding a significant rescue of nuclear H_2_O_2_ levels following CHK1 inhibition (**Figures 4G-H, S7J**). Importantly, DOXY treatment reduced total mitochondrial H_2_O_2_ levels following CHK1i treatment (**Figure S7I**). Finally, we observed a significant decrease in CHK1i-mediated H2AX•S139 phosphorylation following DOXY treatment or SSBP1 depletion (**Figures S7K-L**), supporting our finding that DNA damage following CHK1 inhibition is, in part, due to dysregulation of mitochondrial translation.

We next asked whether SSBP1 could alter nuclear H_2_O_2_ levels following treatment with 9 additional cytotoxic agents which are used in the clinic, finding that SSBP1 depletion substantially decreased nuclear H_2_O_2_ levels upon treatment with ATO, LAP and CPT (**Figure 4I**). Interestingly, cells depleted of SSBP1 were partially protected from any agent that raised nuclear H_2_O_2_ levels (**Figures 4J, S8A**). Because inhibition of GSH biosynthesis increased sensitivity to AUR treatment (**Figures S5D,F**), we wondered whether this GSH-dependency would extend to other agents that increase nuclear H_2_O_2_ levels. Indeed, we find that depletion of glutathione increased the cytotoxicity of AUR, LAP and CPT – all chemotherapies that increase nuclear H_2_O_2_. In contrast, GSH depletion did not affect chemotherapies such as 5FU or 2ME that do not alter nuclear H_2_O_2_ (**Figure S8B**). Mechanistically, we find that treatment with chemotherapies that result in higher levels of nuclear H_2_O_2_ have an increased GSSG:GSH ratio, whereas compounds such as 5FU or 2ME did not alter this ratio (**Figure S8C**). These results suggest the presence of an ‘AND gate’ required to regulate nuclear ROS by chemotherapies: CHK1/SSBP1 control mitochondrial H_2_O_2_ and GSSG:GSH ratio is regulated by chemotherapies. Together, they work in concert to permit mitochondrial H_2_O_2_ to travel to the nucleus, raising peroxide levels in this compartment.

### Depletion of SSBP1 mediates resistance to cisplatin cytotoxicity

Platinum-based chemotherapies are common adjuvant treatments for women with high grade serous ovarian cancers (HGSOCs). When we stratified patients based on SSBP1 mRNA levels, we found that patients with lower levels of SSBP1 transcripts had a shorter duration to tumor recurrence following platinum-based chemotherapy than patients with higher levels of SSBP1 (**Figures S8D-E**). In a cohort of HGSOC patients with differing platinum free intervals (PFIs) we queried SSBP1 protein expression in corresponding tumors, finding lower levels of SSBP1 correlated with a shorter PFI (**Figure 4K**). To probe the role of ROS in cisplatin cytotoxicity, we treated nine ovarian cancer models of different etiologies with NAC, finding a decrease in nuclear H_2_O_2_ and a corresponding increase in the IC_50_ of this drug (**Figures S8F-H**). As we observed in K562 cells, loss of SSPB1 decreased nuclear H_2_O_2_ levels in these models and increased proliferation over a range of cisplatin concentrations (**Figures 4L**, **S8I-J**), suggesting that loss of SSBP1 may promote platinum-based resistance.

Here, we define a nuclear-to-mitochondria ROS sensing circuit that illustrates how nuclear H_2_O_2_ sensing controls mitochondrial translation which in turn regulates the steady state-levels of nuclear ROS, revealing an unexpected connection between DNA damage sensing compartmentalized ROS regulation and resistance to chemotherapy treatment in ovarian cancers (**Figure 4M**).

## Discussion

The majority of cancer patients succumb to disease following the onset of chemoresistance, which we now appreciate arises through both genetic and non-genetic mechanisms (Hanahan and Weinberg, 2011; Senthebane et al., 2017; Vasan et al., 2019; Yeldag et al., 2018; Zheng, 2017). While proteins have long been appreciated as targets of ROS, the identity and functional significance of these targets following chemotherapy is not established. This not only limits our understanding of chemoresistance and normal tissue cytotoxicity, but hampers expanded use of these drugs in the clinic. Herein, using cysteine-focused chemical proteomics and functional genomics, we generated a global portrait of targets for 11 chemotherapies–information that is necessary to understand the cellular response to these drugs. Using this approach, we uncovered a nucleus-to-mitochondria ROS sensing pathway that may play a role in chemoresistance.

Our results suggest that mitochondrial translation is a major determinant of nuclear H_2_O_2_ levels and demonstrate how some chemotherapies collaborate with mitochondrial H_2_O_2_ to increase nuclear H_2_O_2_ and DNA damage. Given that high ROS levels damage nucleic acids (Chao et al., 2013; Juan et al., 2021) it is perhaps not surprising that the nucleus has evolved its own pathway to dynamically respond to this stressor. Crosstalk between these two organelles has been extensively studied in the context of anterograde signaling and corresponding transcriptional regulation (Guaragnella et al., 2018; Lechuga-Vieco et al., 2021). However, our findings provide a direct posttranslational mechanism by which the nucleus leverages the DDR pathway to respond to high levels of nuclear H_2_O_2_ through the concomitant downregulation of mitochondrial translation. The canonical framework for ROS-based activation of DDR relies on direct DNA damage (Ciccia and Elledge, 2010). Our finding of a H_2_O_2_-sensing cysteine within CHK1 that is sufficient for activation of this kinase represents a parallel mechanism by which the cell can respond to changes in altered nuclear H_2_O_2_ levels to protect DNA prior to the accrual of damage. Our cellular and in vitro characterization of nuclear H_2_O_2_ indicate that this ROS is both necessary and sufficient to oxidize C408. Whether other species of ROS or electrophilic compounds can modify C408 remains an open question; however, the conservation of C408 in CHK1 suggests that this residue is likely to be important in multiple kingdoms of life and may function as a general nuclear ROS sensor. Given the role of ROS in DNA damage, the downregulation of mitochondrial translation and resultant decrease in H_2_O_2_ provides a safety check mechanism to preserve genomic integrity. Our findings reveal how the DDR pathway can stymie mitochondrial H_2_O_2_ production, saving the nucleus from ROS-induced damage through the cytosolic retention of SSBP1. How SSBP1 is excluded from the mitochondria remains to be determined but could center on blocking the cleavage of its mitochondrial targeting sequence, which is required to keep SSBP1 in the mitochondria (Gustafson et al., 2019).

More broadly, our findings demonstrate that a decrease in mitochondrial translation leads to a sharp decrease in nuclear H_2_O_2_ levels, suggest that decreasing mitochondrial translation by regulating SSBP1 levels, may be a general mechanism of resistance to chemotherapies that increase steady-state levels of nuclear H_2_O_2_ levels. Indeed, we find that HGSOC patients with lower levels of tumoral SSBP1 have shorter platinum-free interval and directly demonstrate that lowering SSBP1 in multiple models of ovarian cancer confers resistance to cisplatin. Thus, our study suggests that co-treatment of platinum-based chemotherapies with CHK1 inhibitors, which restore SSBP1 localization to the mitochondria and increase ROS, may be an approach to overcome platinum resistance. Moreover, they suggest a general mechanism of resistance for therapeutic agents whose cytotoxicity is mediated in part by increasing the steady-state levels of nuclear H_2_O_2_ and is in accordance with the growing body of evidence that mitochondrial function is a critical determinant of therapeutic response (Fu et al., 2021; Gentric et al., 2019; Shen et al., 2021; Xie et al., 2021).

By combining functional genomics with chemical proteomics, we have defined the mechanisms of sensitivity and resistance to AUR, an emerging anti-cancer drug, that has been at the forefront of efforts to interrogate and interpret ROS biology. Our findings that AUR is sufficient to regulate nuclear H_2_O_2_ levels underscores the importance of its canonical targets TXNRD1/2 in the antioxidant response. Given the high levels of nuclear H_2_O_2_ following AUR treatment, it is perhaps surprising that depletion of one protein, SSBP1, can dramatically decrease H_2_O_2_ at this organelle and corresponding cytotoxicity. Thus, it illustrates the central role of mitochondrial translation in controlling ROS levels at other organelles and its centrality in the response to chemotherapies.

Our previous studies have indicated a strong enrichment for redox-sensitive cysteines as targets of covalent inhibitors (Bar-Peled et al., 2017). Thus, ROS targets defined as essential by functional genomic studies may offer an attractive starting point for the future development of powerful anti-cancer agents that will be more specific than broadly cytotoxic chemotherapies studied herein.

## Acknowledgements

We thank Thomas Michel for providing DAAO-HyPer7 plasmid. We thank all members of the Bar-Peled Lab, David Sabatini and Lee Zou for helpful suggestions. This work was supported by the Damon Runyon Cancer Research Foundation (62-20), the American Association for Cancer Research (19-20-45-BARP), and the American Cancer Society, the Melanoma Research Alliance, the Ludwig Cancer Center of Harvard Medical School, Lungevity, ALK Positive, V-Foundation, Mary Kay Foundation, Paula and Rodger Riney Foundation, and the NIH/NCI (1R21CA226082-01, R37CA260062).

## Author Contributions

J.Z. and L.B-P. conceived and designed the study. J.Z. performed most of experiments with assistance of J.B., J.F., J.O.S., T.W-S., S.H., M.H., T.Y., M.R., H.P., A.E.S., A.D.C., A.H.G., L.S., T.Y.W., B.R.D., N.J.C., T.V., A.D.H., N.W., V.P., Y.M., S.K., E.K. A.P.P., C.M.S. M.P.S., and S.A.B. performed proteomics analysis and interpreted the data. C.M.S., H.B.C., K.W. and M.L. performed bioinformatics analysis. J.Z. and L.B-P. wrote the manuscript with assistance from all the coauthors. L.B-P. supervised the studies.

## STAR+METHODS

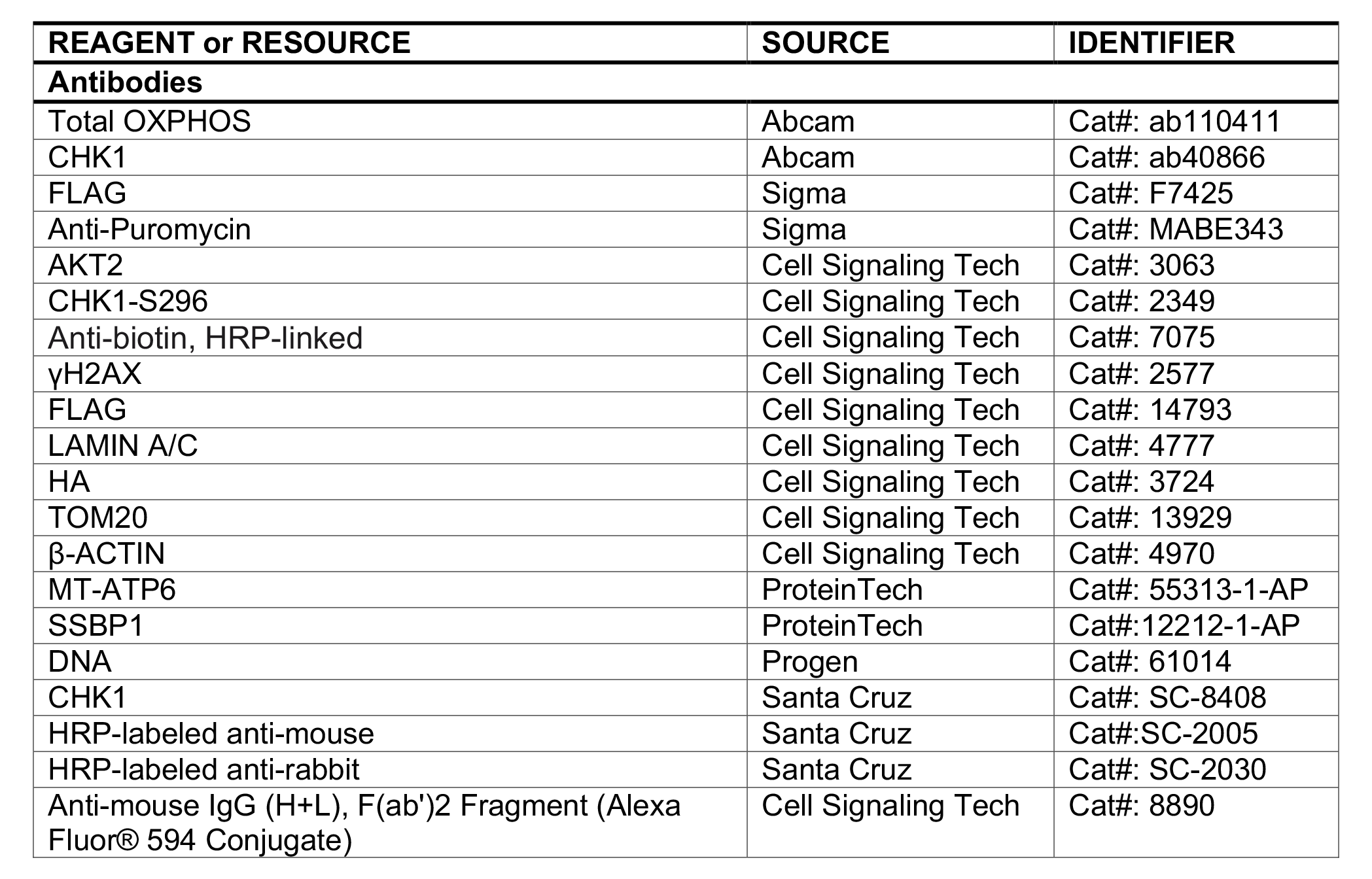

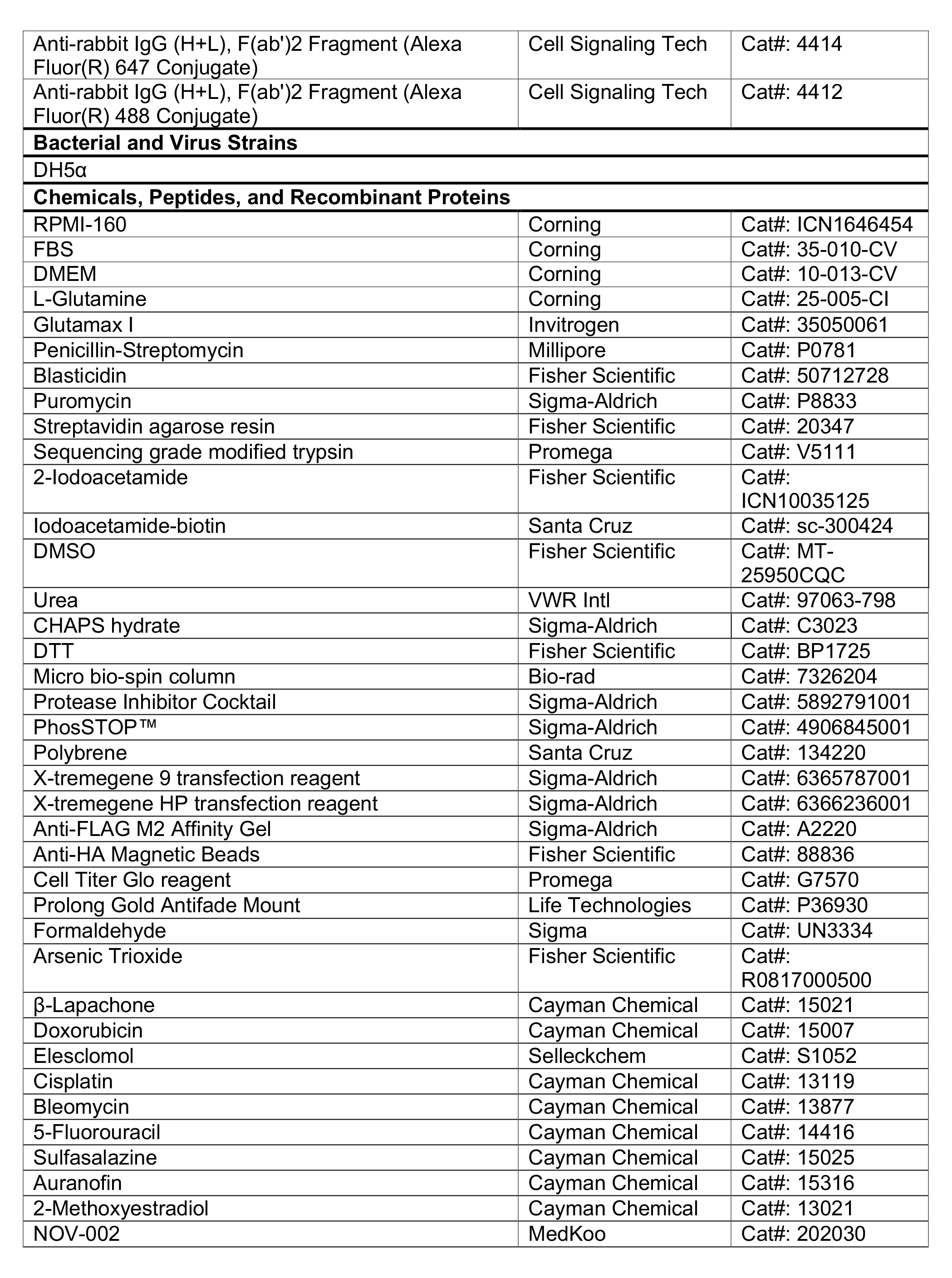

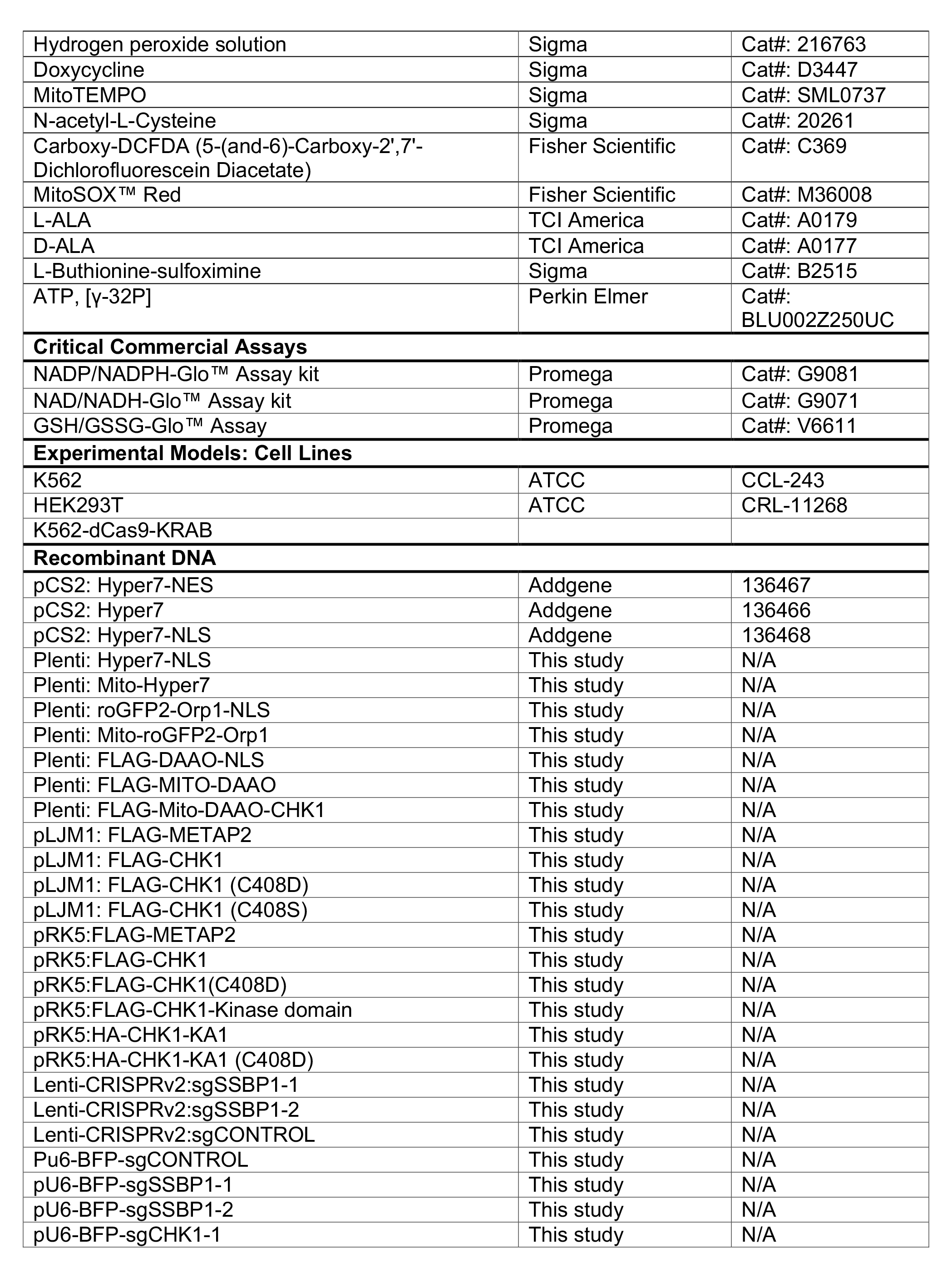

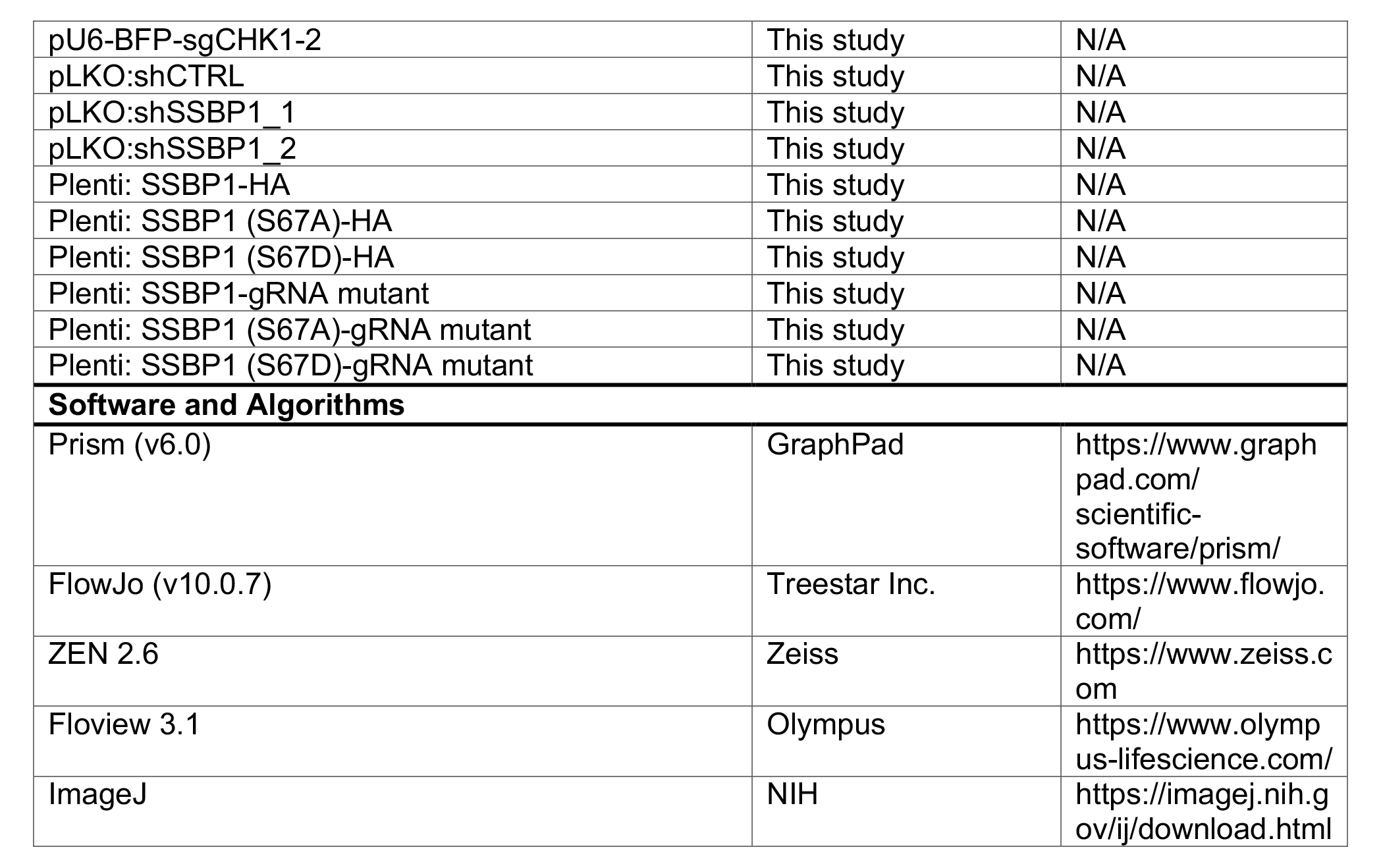
KEY RESOURCES TABLE.

## CONTACT FOR REAGENT AND RESOURCE SHARING

Further information and requests for reagents should be directed to the Lead Contact, Liron Bar-Peled.

## EXPERIMENTAL MODEL AND SUBJECT DETAILS

### Cell lines

All cells were maintained at 37°C with 5% CO2. HEK293T were grown in DMEM (Corning) supplemented with 10% fetal bovine serum (FBS, Corning), Penicillin-Streptomycin (100 mg/ml, Millipore) and L-glutamine (2 mM, Corning). K562 and K562-dCas9-KRAB were grown in RPMI- 1640 (Invitrogen) supplemented with 10% fetal bovine serum (FBS, Corning), Penicillin-Streptomycin (100 mg/ml, Millipore) and 1% GlutaMax (Millipore). All cell lines were routinely tested for Mycoplasma and if not noted elsewhere were obtained from American Tissue Type Collection (ATCC). Whenever thawed, cells were passaged at least three times before being used in experiments.

### cDNA cloning and mutagenesis

cDNAs were amplified using Q5 High-Fidelity 2X master mix (NEB) and subcloned into the pRK5 (Addgene), pLJM1 (Addgene) or pLenti CMV (Addgene) by T4 ligation or Gibson cloning. Site directed mutants were generated using QuikChange XLII site-directed mutagenesis (Agilent), using primers containing the desired mutations. All constructs were verified by DNA sequencing.

### H2DCFDA and MitoSox measurements

For H2DCFDA (Thermo) staining in K562 cells, cells were treated as indicated in the text and were washed with prewarmed PBS and harvested by centrifugation at 1200 *g* at room temperature for 2 mins. The cell pellet was resuspended in PBS with 1 µM of CM-H2DCFDA and incubated for 45 min in a 37°C incubator with controlled CO2 levels (5%). Cells were subsequently washed with PBS. Changes in CM-H2DCFDA fluorescence were determined via flow cytometry using Aurora (Cytek) or CytoFLEX (Beckman Coulter). Data was analyzed using Flowjo v10.6 for FITC intensity. For MitoSox RED (Thermo) staining in K562 cells, 0.5 µM MitoSox Red was added directly to the culture medium and incubated for 20 min in a 37°C incubator with controlled CO_2_ levels (5%). Changes in MitoSox fluorescence were determined by flow cytometry using Aurora or CytoFLEX. PE intensity was analyzed using Flowjo v10.6.

### GSSG/GSH, NADH/NAD+ and NADPH/NADP+ measurement

K562 cells were treated with different chemotherapy drugs in 6-well plates per the timepoints indicated in the text. The ratio of GSSG/GSH, NADH/NAD+ and NADPH/NADP+ was determined using the GSH/GSSG-Glo™ Assay Kit (Promega), NAD/NADH-Glo™ Assay Kit (Promega) and NADP/NADPH-Glo™ Assay Kit (Promega), respectively, following the manufacturer’s protocol. Absorbance was measured using a SpectraMax M5 plate reader (Molecular Devices).

### Confocal imaging of cell lines expressing HyPer7/roGFP2-Orp1 reporters

K562 or K562-dCas9-KRAB cells expressing the indicated HyPer7/roGFP2-Orp1 reporters with a concentration of approximately 5×10^5^ cells/ml were seeded on poly-lysine coated glass bottom dish (fisher) and treated with compounds as indicated in the text. Dishes were firmly mounted on the stage adaptor of the Zeiss 710 Laser Scanning Confocal microscope (Carl Zeiss Inc.). Constant temperature (37 °C), humidity, and 5% CO2 atmosphere were maintained throughout the duration of cell imaging. Images were acquired using a 63X oil objective. The oxidized form of the HyPer7 reporter was detected by exciting HyPer7 expressing cells with a 488-nm laser and measuring emission in the 500-520 nm range. Reduced HyPer7 was detected by exciting HyPer7 expressing cells with a 405-nm laser and measuring emission in the 500-545 nm range. The oxidized form of the roGFP2-Orp1 reporter was measured by exciting roGFP2-Orp1 expressing cells with a 405-nm laser and measuring emission in the 500-520 nm range. The reduced form was measured by exciting roGFP2-Orp1 expressing cells with a 488-nm laser and measuring emission in the 500-545 nm range. Acquisition parameters were identical between samples. Images were processed using the ZEN 2.6 Image software (Carl Zeiss Inc.). Ratiometric images of HyPer7 were processed using ImageJ (NIH). Threshold images after subtraction of background were split into two different channels, divided with Image Calculator. 32-bit ratiometric images were generated and presented in the 16 color mode using Lookup Tables.

### Flow cytometry analysis of cell lines expressing HyPer7/ roGFP2-Orp1 reporters

1.6×10^4^ K562 or K562-dCas9-KRAB cells expressing the indicated HyPer7/ roGFP2-Orp1 reporters were seeded in a 96-well plate for 24 hrs and treated as indicated in the text. HyPer7/ roGFP2-Orp1 oxidation and reduction was determined by flow cytometry using an Aurora (Cytek) or CytoFLEX (Beckman Coulter) measuring emission at 530 nm following excitation at 405 nm or 488 nm. The ratio of λex = 488 nm/ λem = 530 nm to λex = 405 nm/ λem = 530 nm signal for HyPer7 and ratio of λex = 405 nm/ λem = 530 nm to λex = 488 nm/ λem = 530 nm signal for roGFP2-Orp1 was determined using Flowjo v10.6.

### Immunofluorescence

1×10^6^ K562 or K562-dCas9-KRAB cells expressing the indicated HyPer7/ roGFP2-Orp1 reporters or SSBP1-HA were fixed with 4% PFA (EMS) for 15 min and resuspend with PBS to a concentration of approximately 5×10^5^ cells/ml. Cells spun onto coverslips using a thermo scientific cytospin cytocentrifuge. The slides were then rinsed with PBS and cells were permeabilized with 0.1% Triton X-100 in PBS for 10 min. The slides were rinsed with PBS and incubated with primary antibodies in 4% BSA overnight at 4°C. Following three PBS washes, the slides were incubated with secondary antibodies conjugated to the Alexa Fluor® 488 and 594 fluorophores (Invitrogen) for 2 hrs at room temperature. The slides were rinsed and mounted on glass slides using ProLong™ gold antifade mount without or with DAPI (Fisher Scientific). For HEK293T stably expressing FLAG-DAAO (nuclear or mitochondria matrix localized), cells were plated on poly-lysine coated glass coverslips in 12-well tissue culture plates. 48 hrs later, the culture media was removed, and cells were fixed with 4% paraformaldehyde (Electron microscopy services) for 15min. Staining was done as above. Cells were imaged on Zeiss LSM 710 laser scanning confocal microscope or Olympus under 63x oil objective. Images were processed using ZEN 2.6 Image software (Carl Zeiss Inc.) and ImageJ (NIH).

### Cell lysis and FLAG-Immunoprecipitations

Cells expressing the indicated proteins were washed once with ice-cold PBS and lysed using a chilled bath sonicator (Q700, QSonica) in Triton IP buffer (1% Triton X-100 (sigma), 5 mM MgCl_2_, 40 mM HEPES pH 7.4, 10 mM KCl) supplemented with protease inhibitors (Roche), phosphatase inhibitors (Roche) and Benzonase (Santa Cruz). Lysates were clarified by centrifugation at 13000rpm for 10 min. Samples were normalized to 1 mg/ml and boiled following the addition of sample buffer. For FLAG immunoprecipitations, anti-FLAG M2 resin (Sigma) was added to the pre-cleared lysates and incubated 3 hrs at 4°C. Following immunoprecipitation, beads were washed once with Triton IP buffer followed by 3 times with Triton IP buffer supplemented with 500mM NaCl. For in vitro binding assay of CHK1 kinase domain with endogenous CHK1, cells expressing FLAG-CHK1 kinase domain was lysed using IP buffer supplemented with 0.3% CHAPS (Sigma) instead of 1 % Triton. Clarified lysate was pretreated with DMSO or 100 uM H_2_O_2_ for 2 hrs and then incubated with anti-FLAG M2 resin for immunoprecipitation. Loading buffer was added to the immunoprecipitated proteins which were subsequently denatured by boiling for 5min. Proteins were resolved by SDS-PAGE and analyzed by immunoblotting.

### IA-DTB labeling of CHK1

K562 cells stably expressing FLAG-CHK1 or FLAG-C408D•CHK1 were treated with H_2_O or 100 µM H_2_O_2_ for 1h and then incubate with 1mM IA-DTB for 1h. Cell samples were washed with ice-cold PBS and lysed using a chilled bath sonicator (Q700, QSonica) in PBS buffer supplemented with Benzonase (SantaCruz). Samples were adjusted to 2 mg/mL. IA-DTB modified proteins were enriched by the addition of streptavidin beads (Thermo) and following 2 hr. incubation, beads were washed twice with 0.1% IGEPAL (Sigma), PBS and Triton IP buffer supplemented with 500mM NaCl. Proteins were then resolved by SDS-PAGE and analyzed by immunoblotting of FLAG-CHK1 or FLAG-C408S•CHK1.

### Synthesis of NO-DTB probe

Synthesis of phenyl 4-((2-(6-((4S,5R)-5-methyl-2-oxoimidazolidin-4- yl)hexanamido)ethyl)carbamoyl)-2-nitrosobenzoate (**NO-DTB**)(El-Faham and Albericio, 2010; Kuljanin et al., 2021; Lo Conte and Carroll, 2012; Shi and Carroll, 2020) is described as follows:

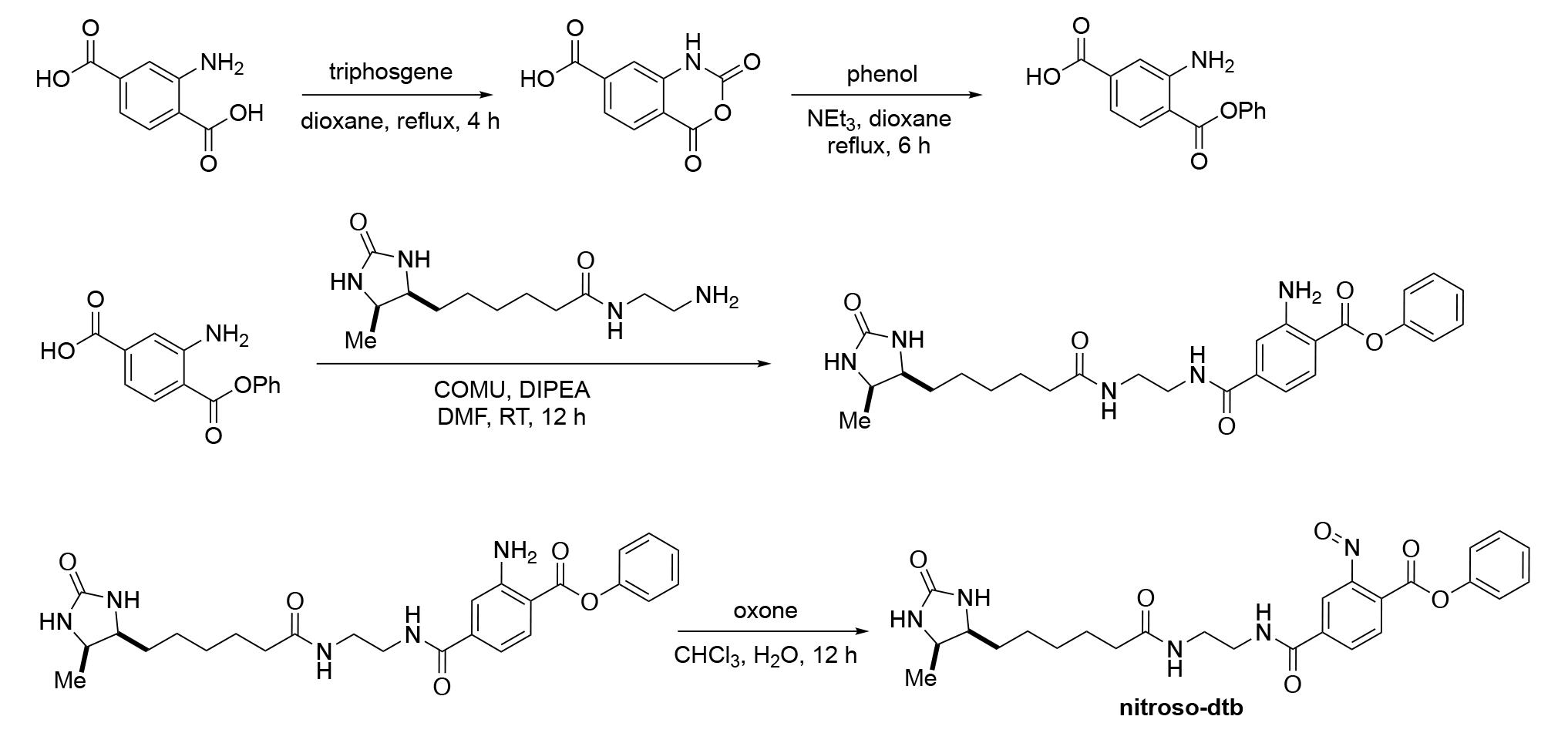

To a solution of 2-aminoterephthalic acid (1.0 g, 5.53 mmol) in 1,4-Dioxane (80 mL) was added triphosgene (1.64 g, 5.53 mmol) at room temperature. The resulting reaction mixture was stirred for 6 hr. at room temperature. The reaction mixture was poured in H_2_O (200 mL) and extracted with ethyl acetate (3 x 75mL). The organic layers were combined, washed with brine and concentrated under reduced pressure to afford 4-carboxylic isatoic anhydride as an off white solid (97%).

Phenol (681 mg, 7.25 mmol) (anhydrous; prior to addition: solubilized in EtOAc, dried with MgSO_4_, concentrated, then placed under vacuum to dry) and triethylamine (1.34 mL, 9.66 mmol) were added to a solution of 4-carboxylic isatoic anhydride (1000 mg, 4.83 mmol) in anhydrous 1,4-dioxane (40 mL). The mixture was stirred under N_2_ and refluxed for 6 h. The crude mixture was concentrated, then diluted with water (15 mL), and the pH was adjusted to 3 with conc. HCl. The solution was extracted with EtOAc (3 x 20 mL) to yield the 3-amino-4-(phenoxycarbonyl)benzoic acid as a bright yellow solid (98%).

To a flame dried 3-necked flask, 3-amino-4-(phenoxycarbonyl)benzoic acid (166 mg, 0.578 mmol), COMU (292 mg, 0.682 mmol), DIPEA (220 mg, 1.70 mmol), and DMF (5 mL) were added. The mixture was stirred under N_2_ for 15 minutes, then N-(2-aminoethyl)-6-((4R,5S)-5- methyl-2-oxoimidazolidin-4-yl) hexanamide (166 mg, 0.625 mmol) was added. The mixture was stirred for 12 hr. at RT, then it was concentrated and purified via column chromatography with DCM/MeOH to afford phenyl 2-amino-4-((2-(6-((4S,5R)-5-methyl-2-oxoimidazolidin-4- yl)hexanamido)ethyl)carbamoyl)benzoate (35%).

A solution of oxone (46.6 mg, 0.303 mmol) in water (3.0 mL) was added to a solution of 1-phenyl2-aminoterephthalate (50.0 mg, 0.101 mmol) in CHCl_3_ (1.0 mL). The reaction was vigorously stirred for 12 hr. The solution was concentrated and purified via column chromatography with DCM/MeOH to afford phenyl 4-((2-(6-((4S,5R)-5-methyl-2-oxoimidazolidin-4-yl)hexanamido)ethyl)carbamoyl)-2-nitrosobenzoate (nitroso-dtb) (21%). 1H NMR (MeOH-d4, 400 MHz, mixture of monomer and dimer 1:1): δ 8.34 (d, J = 8.33 Hz, 0.5H), 8.17-8.04 (m, 2H), 7.63 (s, 1H), 7.49-7.16 (m, 6H), 3.77-3.42 (m, 6H), 2.18 (t, J = 2.18 Hz, 2H), 1.60-1.19 (m, 10H), 1.03 (d, J = 1.03 Hz, 3H). 13C NMR (MeOH-d4, 100 MHz, mixture of monomer and dimer): δ 175.5, 150.9, 133.4, 130.0, 129.4, 129.2, 126.2, 121.8, 121.3, 113.77, 113.11, 56.0, 51.3, 38.45, 35.7, 29.3, 28.9, 25.8, 25.5, 14.27. ESI-LCMS calcd. for C_26_H_31_N_5_O_6_ (M-H) 510.2, found 510.2.

**Figure A.**
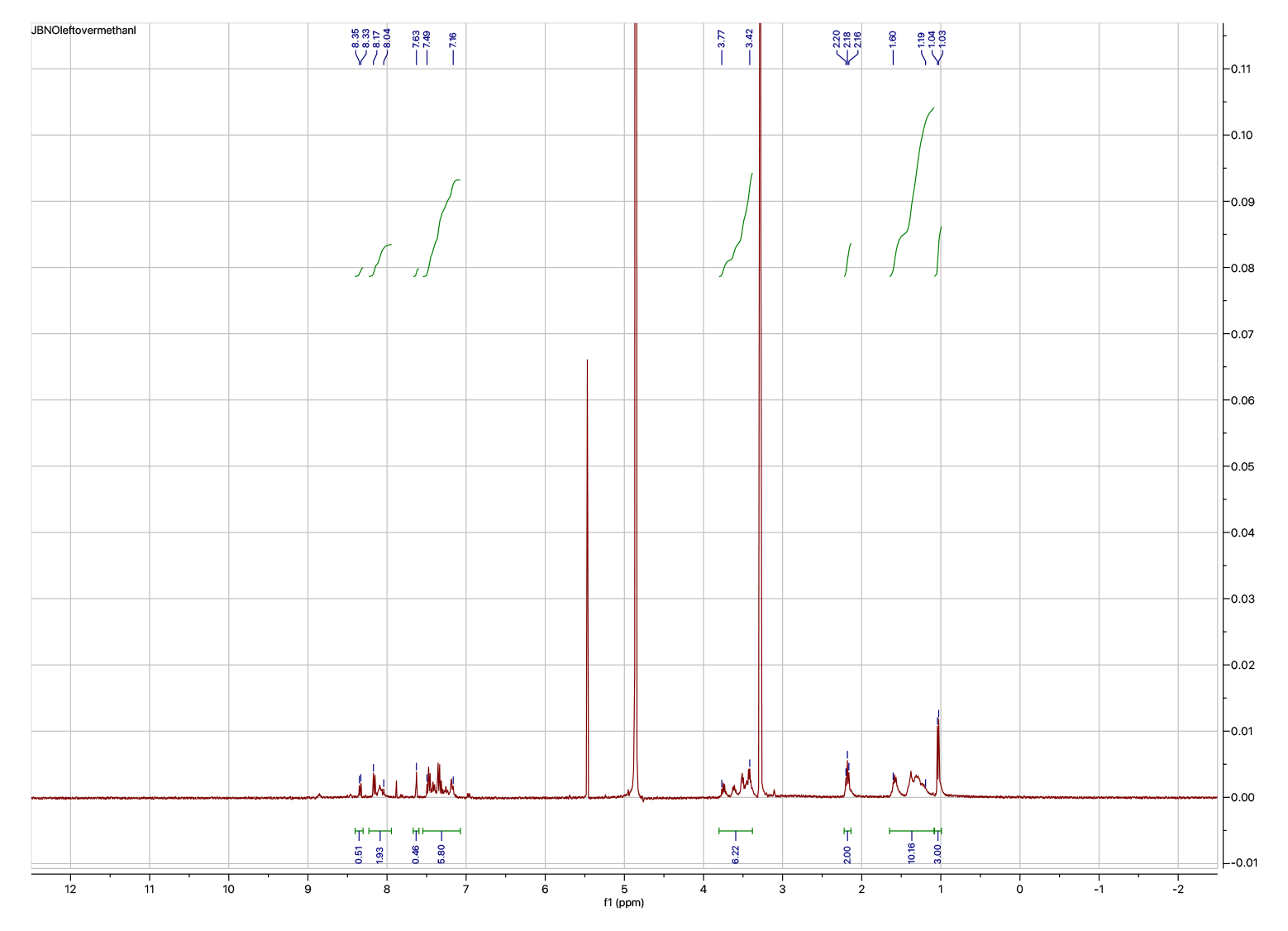
^1^H NMR of Nitroso-DTB.

**Figure B.**
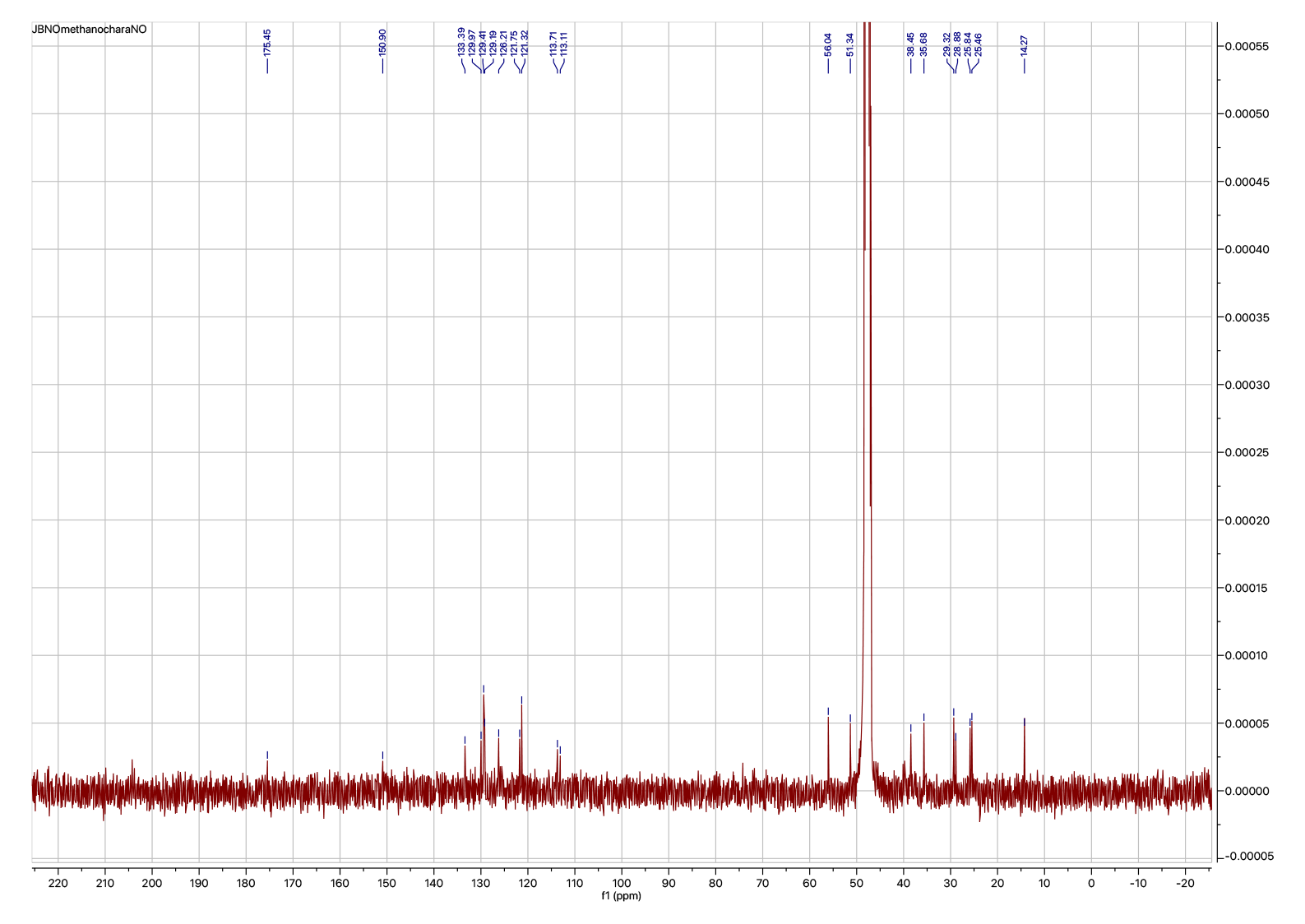
^13^C NMR of Nitroso-DTB.

### Detection of CHK1 Sulfinylation

Detection of CHK1 Sulfinylation by NO-DTB labeling was performed as previously reported (Lo Conte et al., 2015; Yang et al., 2016). In brief, K562 cells stably expressing FLAG-CHK1 or FLAG- C408S•CHK1 were treated with H_2_O or 100 µM H_2_O_2_ for 1hr as above. Cell samples were washed with ice-cold PBS and lysed using a chilled bath sonicator (Q700, QSonica) in Triton IP buffer (1% Triton X-100 (sigma), 5 mM MgCl2, 40 mM HEPES pH 7.4, 10 mM KCl) supplemented with protease inhibitors (Roche), phosphatase inhibitors (Roche), Benzonase (Santa Cruz) and 5mM DTT. Lysates were clarified by centrifugation at 13000rpm for 10 min. Free thiols were trapped by incubation with 2 mM of 4,4’-dithiodipyridine (4-DPS) at room temperature for 1 hr. and subsequently the buffer was exchanged using one Micro Bio-Spin column pre-equilibrated with 100 mM HEPES, pH 8.5, 100 mM NaCl. DPS-free lysates were then reacted with 500 µM NO-DTB in the dark at room temperature with rotation for 1 hr. Total protein was purified by the addition of a chloroform-methanol solution (4:4:1, methanol, water, chloroform) to each sample and precipitated following centrifugation at 4200 RPM for 10 min. The protein disc was isolated, washed once in methanol, resuspended in Buffer X1 (9 M Urea, 10 mM DTT, 50 mM tetramethylammonium bicarbonate (TEAB) and incubated for 20 min at 65°C. NO-DTB modified proteins were then enriched by the addition of streptavidin beads (Thermo) and following 2 hr. incubation, beads were washed twice with 0.1% IGEPAL (Sigma), PBS and Triton IP buffer supplemented with 500mM NaCl. Proteins were then resolved by SDS-PAGE and analyzed by immunoblotting of FLAG-CHK1 or FLAG-C408S•CHK1.

### In vitro CHK1 kinase assay

K562 cells stably expressing FLAG-CHK1 or FLAG-C408D•CHK1 were lysed by sonication in Triton IP buffer and immunoprecipitated using anti-FLAG M2 beads as described above. Immobilized FLAG-CHK1 was dephosphorylated by treating the protein with calf alkaline phosphatase (NEB) for 1 hr. at 37°C. Immobilized FLAG-CHK1 was subsequently washed 3 times in CHK1 kinase buffer (10 mM HEPES, pH 7.5, 10 mM MgCl_2_, 10 mM MnCl_2_) and the kinase assay was initiated by adding 1 mM ATP (Sigma) and the indicated compounds and incubating at 37°C for 45 min. The reaction was stopped by washing the samples once with ice-cold CHK1 kinase buffer and adding loading buffer. Proteins were resolved by SDS-PAGE and analyzed by immunoblotting. For in vitro phosphorylation of SSBP1 by CHK1, HEK293T cells transiently expressed SSBP1-HA or SSBP1 (S67A)-HA were lysed by sonication in Triton IP buffer and immunoprecipitated using anti-HA magnetic beads. Immobilized SSBP1-HA was dephosphorylated by treating the protein with calf alkaline phosphatase (NEB) for 1 hr. as above. Immobilized SSBP1-HA or SSBP1 (S67A)-HA was subsequently incubated with HA-CHK1 or HA- METAP2 cell lysate and washed 3 times in CHK1 kinase buffer (10 mM HEPES, pH 7.5, 10 mM MgCl_2_, 10 mM MnCl_2_). The kinase assay was initiated by adding 100 μM ATP (Sigma), 2 μCi [γ- 32P] ATP (PerkinElmer), incubated at 30°C for 30 min, and then boiled at 95°C for 5 min in 5X sample buffer. Phosphorylation was assessed by 16% SDS-PAGE and Coomassie Brilliant Blue staining followed by autoradiography.

### Mitochondria translation measurement

K562 cells were treated with 2.3µM Doxycycline to inhibit mitochondrial translation. After 72 hrs treatment, the cells were harvested and refreshed with RPMI medium with DMSO or 2 µM CHK1i. Cell samples were harvested at different timepoint as indicated in the text and lysed as above. Proteins were then resolved by SDS-PAGE and analyzed by immunoblotting of MT-COXII.

### isoTOP-TMT sample preparation

isoTMT samples were prepared as described in(Bar-Peled et al., 2017), with the modifications noted below. Briefly, K562-dCas9-KRAB cells were treated at 37°C with the indicated compounds for the noted time in the text. Cells were harvested by centrifugation at 1200 *g* for 2 min and then washed once with ice-cold PBS and lysed in PBS with Benzonase (Santa Cruz) by using a chilled bath sonicator. Samples were clarified by centrifugation for 3 min at 300 *g*. Samples were adjusted to 2 mg/mL and incubated with 100 µM of iodoacetamide-desthiobiotin (IA-DTB, Santa cruz) for 1 hr. at room temperature. Alkylation was terminated by the addition of a chloroform-methanol solution (4:4:1, methanol, water, chloroform) to each sample and proteins were precipitated following centrifugation at 4200 RPM for 10 min. The protein disc was isolated, washed once in methanol, resuspended in Buffer X1 (9 M Urea, 10 mM DTT, 50 mM tetramethylammonium bicarbonate (TEAB) and incubated for 20 min at 65°C. Samples were subsequently alkylated with 500 mM Iodoacetamide for 30 min at 37°C and digested for 3 hrs with Trypsin (Promega). IA-DTB modified peptides were enriched by the addition of streptavidin beads (Thermo) and following 2 hr. incubation, beads were washed twice with 0.1% IGEPAL (Sigma), PBS and H_2_O. Peptides were eluted with a mixture of 50:50:0.1 (Acetonitrile: H_2_O: Formic Acid) and subsequently dried.

### TMT-Labeling

Samples were prepared as previously described(Possemato et al., 2017). Briefly, 20µg of peptides from each sample were labeled with isobaric tandem-mass-tag (TMT) reagents (Thermo Fisher Scientific, San Jose, CA) in 20 mM pH 8.5 HEPES with 30% acetonitrile (v/v) with 50 ug of TMT reagent. The reaction was quenched for 15 min by adding hydroxylamine to a final concentration of 0.3% (v/v). Samples were combined, dried, purified over SepPak C18 columns, and dried again. Samples were then resuspended in 40 μL of basic reverse phase (bRP) buffer A (10 mM NH4HCO2, pH10, 5% ACN) and separated on a Zorbax Extended C18 column (2.1 × 150 mm, 3.5 µm, no. 763750-902, Agilent) using a gradient of 10-40% bRP buffer B (10 mM NH_4_HCO_2_, pH 10, 90% ACN). 96 fractions were collected before concatenation to 12 or 24 fractions. Each fraction was dried and desalted over a C18 STAGE-Tip prior to analysis by mass spectrometry.

### LC–MS Analysis of Total Protein Fractions

Samples were analyzed on an Orbitrap Fusion Lumos mass spectrometer (Thermo Fisher Scientific, San Jose, CA) coupled with a Proxeon EASY-nLC 1200 liquid chromatography (LC) pump (Thermo Fisher Scientific, San Jose, CA). Peptides were separated on a 100 μm inner diameter microcapillary column packed with ∼40 cm of Accucore150 resin (2.6 μm, 150 Å, Thermo Fisher Scientific, San Jose, CA). For each analysis, we loaded approximately 1 μg onto the column. Peptides were separated using a 2.5 hr. gradient of 6–30% acetonitrile in 0.125% formic acid with a flow rate of 550 nL/min. Each analysis used an SPS-MS3-based TMT method ((McAlister et al., 2014; Paulo et al., 2016; Ting et al., 2011)), which has been shown to reduce ion interference compared to MS2 quantification. The scan sequence began with an MS1 spectrum (Orbitrap analysis, resolution 120,000; 350-1400 m/z, automatic gain control (AGC) target 4.0 × 105, maximum injection time 50 ms). Precursors for MS2/MS3 analysis were selected using a Top10 method. MS2 analysis consisted of collision-induced dissociation (quadrupole ion trap; AGC 2.0 × 104; normalized collision energy (NCE) 35; maximum injection time 120 ms). Following acquisition of each MS2 spectrum, we collected an MS3 spectrum a method in which multiple MS2 fragment ions are captured in the MS3 precursor population using isolation waveforms with multiple frequency notches. MS3 precursors were fragmented by HCD and analyzed using the Orbitrap (NCE 65, AGC 3.5 × 105, maximum injection time 150 ms, isolation window 1.2 Th, resolution was 50,000 at 200 Th).

### MS Data Processing and Analysis

MS spectra were evaluated using Comet and the GFY-Core platform (Harvard University)(Eng et al., 2013; Eng et al., 1994; Huttlin et al., 2010; Villen et al., 2007). Searches were performed against the most recent update of the Uniprot Homo sapiens database with a mass accuracy of +/-50ppm for precursor ions and 0.02 Da for product ions. Static modification of lysine and N-termini with TMT (229.1629 Da) and carbamidomethylation (57.0215 Da) of cysteine were allowed, along with oxidation (15.9949 Da) of methionine residues and modification (398.2529 Da) of cysteine residues as variable modifications. Results were filtered to a 1% peptide-level FDR with mass accuracy +/-5ppm on precursor ions and presence of a modified cysteine residue for Cys-Mod samples. Results were further filtered to a 1% protein level false discovery rate. TMT quantitative results were generated in GFY-Core. For TMT-based reporter ion quantitation, we extracted the summed signal-to-noise (S/N) ratio for each TMT channel and found the closest matching centroid to the expected mass of the TMT reporter ion. MS3 spectra with TMT reporter ion summed signal-to-noise ratios less than 100 were excluded from quantitation.

### Ratio and Median Calculation

Abundances for each peptide corresponding to a site on a canonical protein were totaled and normalized to the median for each agent and control. The ratio (R) between the abundances for the control and agent was calculated. For each protein and agent, the median was calculated among the ratios for each site on the protein if there were 3 or more sites identified.

### Circos plot

The R package “circlize” was used to create a chord diagram where each point around the circle represented a site with R > 1.5 in a particular treatment and edges were drawn between any sites shared between two treatments (https://cran.r-project.org/web/packages/circlize/citation.html).

### Clustering Analysis

The UMAP embedding was calculated with the umap.UMAP function of the Python umap.umap_ package (n_neighbors = 15 and random_state = 42). Reactive cysteines were clustered using K-means clustering.

### Structural Analysis

We downloaded PDB files mapping to 4199/15165 commonly detected cysteines (**Supplementary Table 9**) and mapped PDB chains to UniProtKB entries (Martin, 2005). Centroids for each amino acid within a structure were computed, and all residues with Euclidean distance ≤ 10Å of a cysteine of interest were included for further analysis. The distances of these neighbors were then rank ordered in ascending fashion, and these sorted lists were used to generate pLogo motifs (O’Shea et al., 2013). The neighbors of cysteines with 0.7 ≤ max(R) ≤ 1.3 were used to calculate background probabilities in the pLogo algorithm. To obtain spatial enrichment of residues near cysteines of interest we binned the 10Å radius around cysteines of interest into 0.5Å intervals (O’Shea et al., 2013). In order to ascertain the concordance between neighbors in the primary sequence (derived from UP000005640_9606.fasta) and 3-dimensional space, we tabulated the frequency of agreement between the nth-nearest neighbor in 3D space and the ±nth-nearest neighbor in linear sequence on all reactive cysteines. All scenes were generated in PyMOL (PyMOL Molecular Graphics System, version 2.5.2, Schrödinger).

### Reactive Cysteine Signature Score

The score was calculated by multiplying the number of cysteines with a ratio above 1.5 for all agents by the median of those ratios above 1.5, only including cysteines which had observations for all agents.

### Protein Localization Analysis

The most common annotation among the five cell lines (A431, H322, HCC827, MCF7, and U251) from SubCellBarCode (SCBC) was calculated for each protein. Annotated localizations from UniProt and Protein Atlas were compared to extract matching terms and prefixes. Terms, prefixes, SCBC neighborhoods were mapped to a discrete list of subcellular localizations, and the consensus localization was calculated from any localization found in two of the three datasets or SCBC if there were no matching localizations between UniProt and Protein Atlas. Additionally, UniProt annotations tagged with “ECO:0000269” (indicating manually curated annotations derived from published experimental evidence) were extracted and mapped to the same list of subcellular localizations(Orre et al., 2019).

### Gene Ontology Analysis

The R package “topGO” was used to perform a gene ontology analysis on the set of proteins containing any cysteine with R > 1.5 for a single chemotherapy. Molecular function, cellular compartment, or biological process terms were derived from Bioconductor’s org.Hs.eg.db database, and enrichments were computed using the classic method and Fisher’s exact test (Alexa et al., 2006).

### Annotations

Ubiquitination sites from PhosphoSitePlus (Hornbeck et al., 2015) were matched to nearby sites identified by mass spectrometry. Essentiality scores were derived from DepMap (Meyers et al., 2017; Tsherniak et al., 2017). Ribosomal proteins were annotated from the Ribosomal Protein Gene Database. Domain annotations are from UniProt. Functional annotations were taken from the Gene Ontology Annotation database(Huntley et al., 2015). BiomaRt (Durinck et al., 2009) was used to map proteins between UniProt, ensembl, entrez, and PDB IDs and gene names.

### Genome-wide CRISPRi screen

The CRISPRi screen in K562 cells was conducted as previously described (Gilbert et al., 2014). Briefly, K562 cells stably expressing dCas9-KRAB were infected with a genome-wide CRISPRi library cloned into the pU6-BFP vector, ensuring a multiplicity of infection ∼0.3 following 3 days puromycin selection. Cells were allowed to recover for 1 day and an initial input was taken with the number of infected cells corresponding to 1000X the size of the library (∼200×10^6^ cells). The screen was initiated by treating 240×10^6^ with DMSO or 1 µM Auranofin, maintaining this cell number and compound for 10 population doublings. At the end of the screen, cells were harvested and genomic DNA was extracted using Macherey Nagel Blood XL kit (Macherey). Libraries were generated from each sample by PCR based amplification of the sgRNA amplicon from 200 µg of genomic DNA using custom PCR primers harboring an index primer and illumina 5’ and 3’ adaptors. Libraries were pooled and analyzed on a NextSeq500 (Illumina) use single end 75bp reads. sgRNAs were mapped and quantified using the Screen Processing pipeline(Gilbert et al., 2014). The enrichment for each sgRNA was calculated by taking the log_2_ ratio of (sgRNA counts, treatment/sgRNA counts input). The CRISPR score for each gene was calculated by subtracting the enrichment score for ARU treated samples from the enrichment score for DMSO treated samples.

### Cluster analysis

To identify functionally related genetic clusters from CRISPR-Cas9 based essentiality screens, we utilized the depmap resource (https://depmap.org/portal/download/; access: 03/23/2020). The Achilles_gene_effect.csv (Version: DepMap Public 20Q1) matrix summarizes essentiality scores obtained from genome-wide loss-of-function screens from a set of 18,333 genes among 741 different cell lines. Gene-gene correlations across cell lines were computed in R using the cor() function (parameters: method “pearson”; use = “complete.obs”) generating a matrix with 18,333 x 18,333 entries representing correlations of each gene pair. We extracted correlations with values > 0.25 and listed them together with its corresponding gene pair. These list entries were considered as edges of an undirected graph and were imported as network in Cytoscape (Version 3.6.0). The gene network was further dissected using the random walk Markov Clustering algorithm within the Cytoscape plugin clusterMaker (parameters: “granularity” = 1.8; “number of iterations” = 16; “input” = edgelist with gene-gene-correlation as weight). This operation created gene groups with 1 gene up to 695 genes with each gene only occurring in 1 group (= cluster_ID). We proceeded with clusters that contained 3 or more genes resulting in 1028 distinct groups containing a total of 10474 genes. Genes were assigned to list entries in R by their cluster_ID.

Competition scores for cysteines of individual peptides from chemical proteomics experiments were transformed in a binary matrix (1 for competition values ≥ 1.5 = “hit” and 0 for < 1.5 = “no hit”) in R. All peptide hits for each condition were collapsed per protein/gene. Hits were summed up for all genes of a cluster (cluster score) and normalized by a cluster size factor ((size cluster of interest/size largest cluster)^2.03). Hits on the cluster level were summarized by summing up hits for each gene within the cluster for each condition.

Clusters were plotted as circles in Cytoscape (= nodes with no edges) and scaled by cluster scores. Proteomics hits of genes in the cluster were depicted as ratios in pie chart superimposed on the cluster node.

### Target Prioritization analysis

For identifying top scoring gene/protein candidates from chemical proteomics and CRISPRi screens, we combined both data sets. All data processing steps were performed in R. The chemical proteomics data matrix was imported and NA containing rows were removed. We collapsed chemical proteomics competition values per protein/gene name by only keeping the top scoring peptide and ignoring others.

CRISPR scores from loss-of-function screens were imported in R and NA containing cells were removed. The chemical proteomics and CRISPPR matrices were merged using merge() in R by the gene/protein name as identifier. We calculated priority scores by multiplying CRISPR scores with proteomics competition values for each gene/protein.

To identify functionally related protein groups, we utilized the STRING resource (https://string-db.org/;access01/08/2022). We considered genes/proteins as hits if they scored with a CRISPR value of < - 0.75 or > + 0.75 and with a proteomics competition value of > 1.5. For these genes/proteins we extracted the 10 top scoring interactors from the STRING network and considered these 11 genes/proteins as a functional group. For each group we averaged priority scores and corrected them with a penalty score of 1% (multiplication with 0.99^(number of missing genes in the group)) per missing gene in the priority score matrix. These optimized priority scores were plotted as nodes in Cytoscape (= nodes with no edges) and scaled by priority scores (color, node size, label size).

### CRISPRi-mediated knockdown in K562 cells

sgRNAs targeting the promoters of CHK1 and SSBP1 were cloned into pU6-BFP (Addgene: #60955). sgRNA-encoding plasmids were co-transfected with pspAX2 envelope and CMV VSV-G packaging plasmids into 1.8X10^6^ HEK293T cells using the Xtremegene 9 transfection reagent (Sigma-Aldrich). Virus-containing supernatants were collected 48 hrs after transfection and used to infect K562- dCas9-KRAB cells in the presence of 10 mg/ml polybrene (Santa Cruz). Twenty-four hours post-infection, fresh media was added to the infected cells which were allowed to recover for an additional twenty-four hours. Puromycin was then added to cells, which were analyzed after 3 days after selection was added.

### Cell viability assay

2500 cells were seeded in 96-well plates per well in 100 µl medium. K562 cells were subsequently treated with the indicated compounds, whereas adherent cells were treated 24 hrs after seeding. 96 hrs after treatment Cell Titer Glo (Promega) was added to each well sample and the luminescence read on the SpectraMax M5 plate reader (Molecular Devices). For viable cell counting using trypan blue, 0.5 ml of a suitable cell suspension (dilute cells in complete medium without serum to an approximate concentration of 1 x 10^5^ to 2 x 10^5^ cells per ml) was mixed with 0.1 ml of 0.4% trypan blue staining and filled in a hemocytometer cell counting.

### Immunohistochemical (IHC) staining

Immunohistochemistry for SSBP1 were conducted using a two-step protocol (GTVisionTMIII). Briefly, TMA sections were washed with phosphate-buffered saline (PBS) after rehydration and then the antigens were retrieved by boiling the TMA slides in citrate buffer (pH 6.2) at 100°C for 10 min. The TMAs were permeabilized with 0.1% Triton-X-100 for 10 min and then treated with 3% hydrogen peroxide for 10 min to block endogenous peroxidase activity. The TMAs were blocked with 6% BSA for 1 h at room temperature (RT) and incubated in a humid chamber with SSBP1 primary antibody (1:50) for 1 hr. Following washing with PBS, all the TMAs were incubated with secondary antibody (HRP-labeled anti-rabbit antibody 1:50) at RT for 1h. The sections were counterstained with hematoxylin and mounted after clearing with xylene. For hematoxylin and eosin staining, TMA sections were washed with phosphate-buffered saline after rehydration and stained for 1min in hematoxylin solution. The slides were then washed in running tap water. After washing, the slides were dipped in Eosin solution for 1 minute and rinsed with absolute alcohol. The TMA slides were dipped two times in Xylene solution for 1 minute each and mounted with the DPX mountant (Sigma). SSBP1 immunohistochemistry were semi-quantitatively scored via light microscopy by a board-certified pathologist (Z.O.S.) using a 3-point index scoring system for relative expression intensity.

### TCGA analysis

RNA-seq count data was downloaded from the TCGA dataset (Cancer Genome Atlas Research, 2011; Villalobos et al., 2018) and parsed for patients that were treated with platinum-based chemotherapies (cisplatin or carboplatin). Patients were censored if clinical notes indicated confounding treatment or inconclusive progression. For each patient normalized SSBP1 RNA-counts were extracted, and patients were separated into an SSBP1-high and SSBP1-low expression cohort. A Kaplan-Meyer curve was generated in PRISM (ver 8.0) comparing time to tumor recurrence as determined in (Villalobos et al., 2018) for each patient cohort.

### Statistical analysis

Statistical analyses were performed with Excel (Microsoft) and Prism (GraphPad). Error bars represent mean ± s.d. Statistical comparisons were analyzed using two-tailed Student’s t-test with P values indicated in figure legends and source data. Data were considered statistically different at P < 0.05. P < 0.05 is indicated with single asterisks, P < 0.001 with double asterisks, and P < 0.0001 with triple asterisks.

(Fang and Zhang, 2020; Gerber et al., 2018; Hecht, 2000; Longley et al., 2003; Miller et al., 2002; Montero et al., 2012; O’Day et al., 2013; Pardee et al., 2002; Qu et al., 2010; Rivankar, 2014; Rosenberg et al., 1969; Shitara et al., 2017; Sweeney et al., 2005; Tevaarwerk et al., 2009; Xie et al., 2015)

(Bindels et al., 2017; Bulina et al., 2006; Idevall-Hagren et al., 2012; Katajisto et al., 2015; Lee et al., 2020; Malinouski et al., 2011; Pan et al., 2017; Ray et al., 2020; Sun et al., 2010)

---

**Figure S1:**
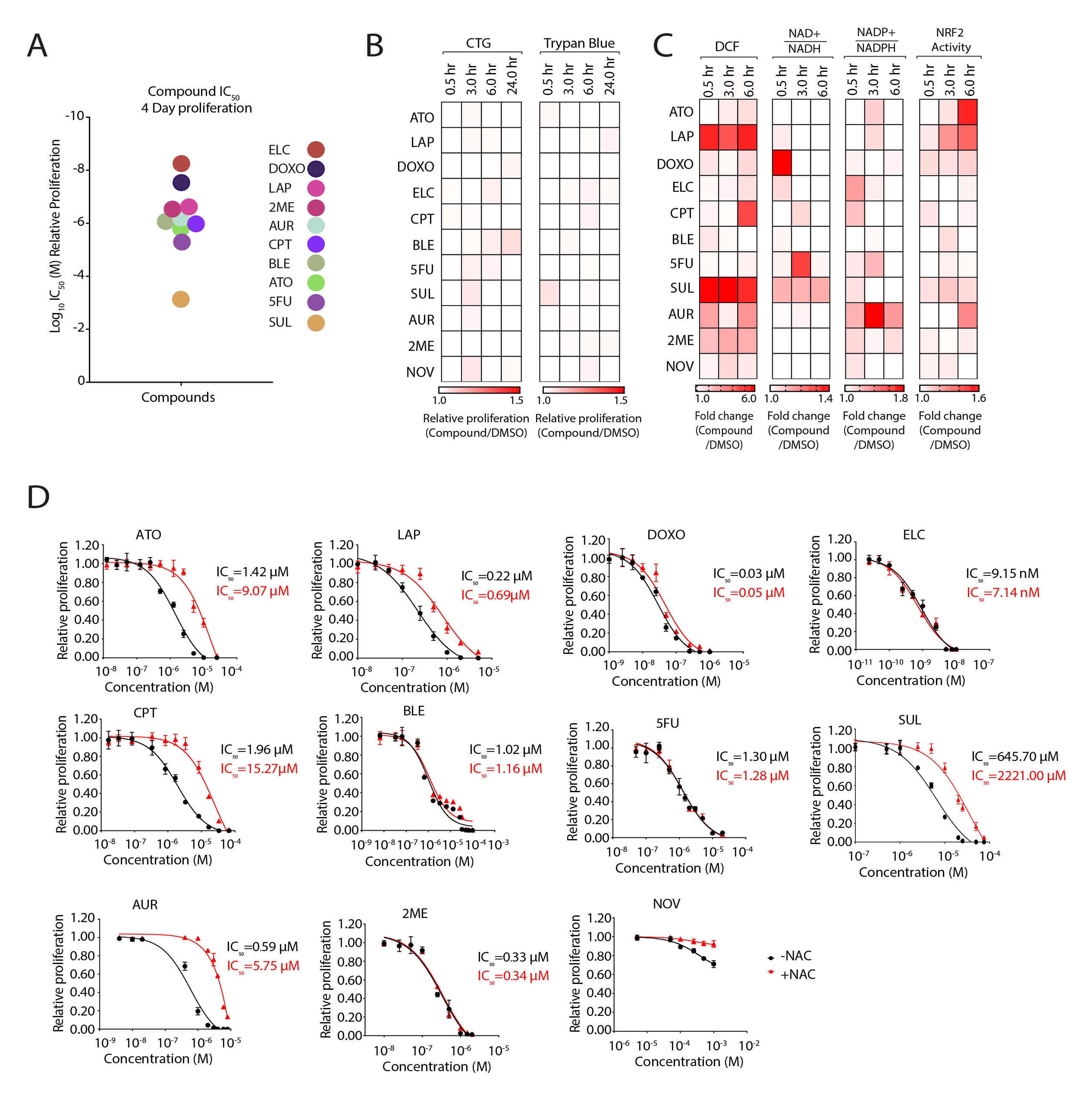
Chemotherapy treatment regulates cellular antioxidant levels. (A) Half-maximal inhibitory effect (IC_50_) of chemotherapies used in this study. Proliferation of K562 was determined after 96 hrs. by measuring cellular ATP concentrations. (B) Short-term treatment with chemotherapies does not alter K562 cell proliferation. Heatmaps depicting changes in K562 proliferation following treatment of the indicated compounds at 5-fold their IC_50_ concentrations as determined in (A). Concentrations of the compounds are: 5.1µM ATO, 1.5µM LAP, 0.1µM DOXO, 30nM ELC, 8.3µM CPT, 7.1µM BLE, 6.0µM 5FU, 0.5mM SUL, 2.5µM AUR, 1µM 2ME or 0.5mM NOV for 0.5, 3, 6 and 24 hrs. Proliferation was determined by measuring cellular ATP concentrations (left) or trypan blue staining (right). (C) Characterization of changes in DCF fluorescence and antioxidant response following chemotherapy treatments. K562 cells treated with the indicated compounds at the noted time points and heatmaps illustrating changes were generated for DCF intensity, NAD+/NADH ratio, NADP+/NADPH ratio, and NRF2 activation. (D) Supplementation with NAC increases the IC_50_ for some chemotherapies. K562 cells were co-treated with the indicated concentrations of each chemotherapy and either 5 mM NAC or vehicle control. Relative proliferation was determined as described in (A).

**Figure S2:**
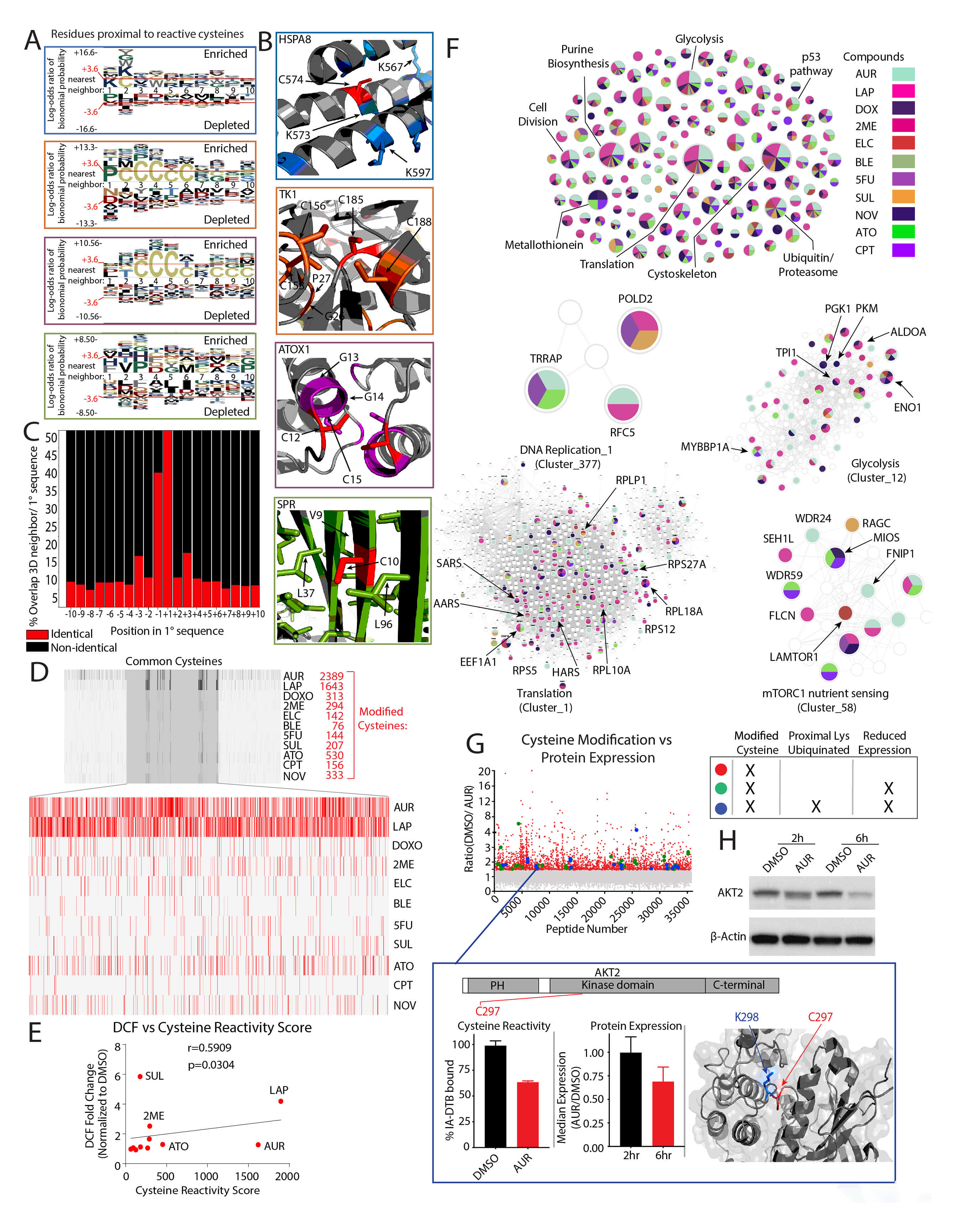
Chemotherapy treatment regulates cysteine reactivity. (A) pLOGO plots of residues found within 10Å of cysteines from different clusters. (B) Examples of cysteines (red) targeted by chemotherapy and their nearest residue neighbors. HSPA8: 1FE0 (Wernimont et al., 2000); TK1:1W4R (Birringer et al., 2005); ATOX1: 4KBQ:(Birringer et al., 2005); SPR:1Z6Z. Enriched amino acids were colored based on corresponding cluster (see also Figure 1B). (C) Bar chart showing disagreement between primary sequence neighbors and 3D neighbors of reactive cysteines. (D) Barcode plot of modified cysteines following treatment with the indicated agents. Inset, a cysteine reactivity score (CRS) is calculated based on 2400+ modified cysteines (R≥1.5) quantified in all treatments (see also **Supplemental Table 2**). (E) Correlation analysis between DCF staining and cysteine reactivity score. (F) Analysis of cysteine reactivity changes following chemotherapy treatment localized to corresponding cellular pathways. Pathways were determined by clustering genes based on co-essentiality. (G) AUR reduces protein expression following cysteine modification. Left, plot of cysteine-reactivity changes in K562 following AUR treatment. Center, expression of AKT2 is decreased following AUR treatment. Right, crystal structure of AKT2 (PDB ID: 3D0E) highlighting the location of C297 and K298. K298 has been previously annotated as ubiquitinated. (H) Immunoblot analysis of AKT2 levels following AUR treatment.

**Figure S3:**
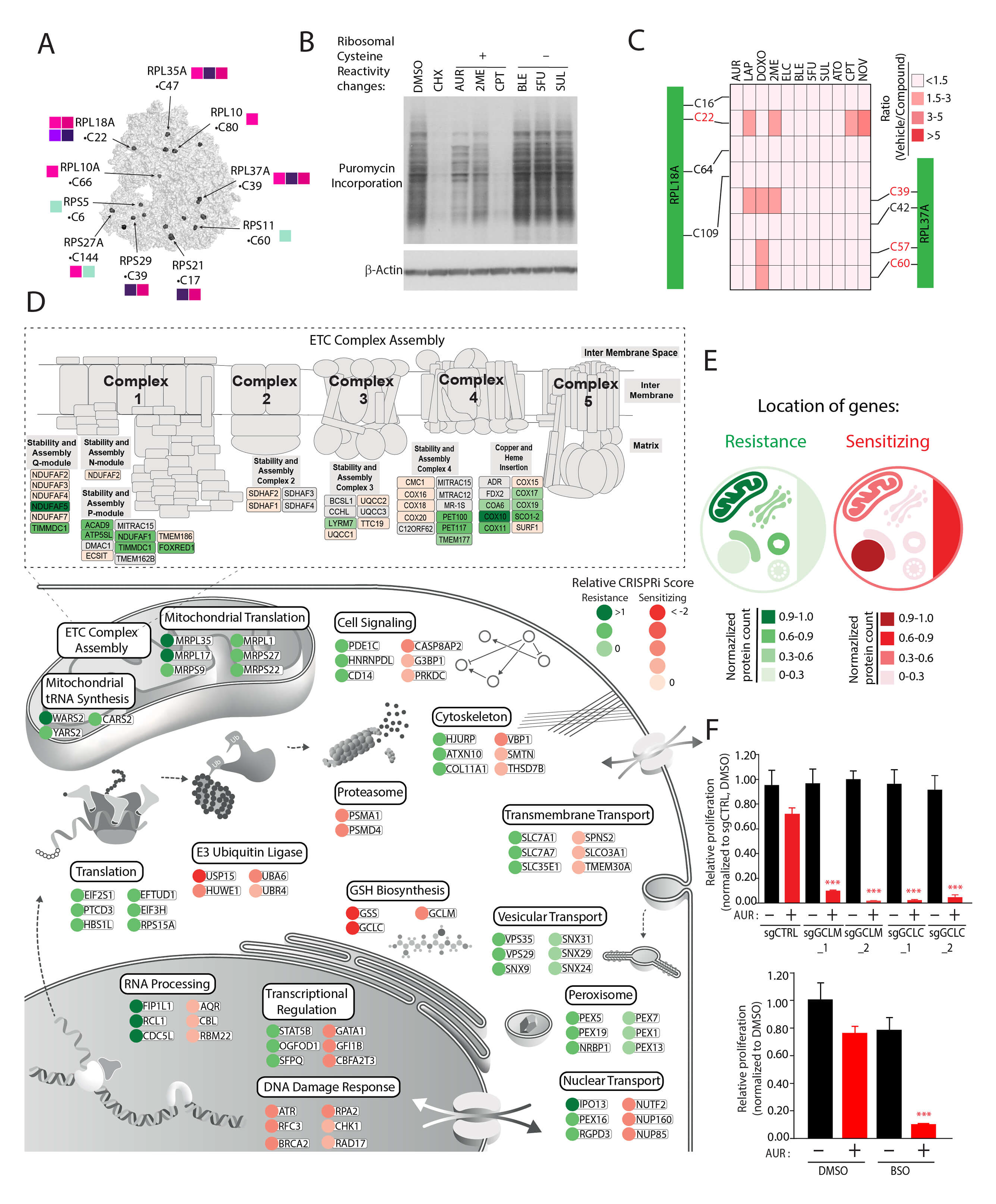
Defining mechanisms of sensitivity and resistance to AUR. (A) Localization of chemotherapy regulated cysteines on the ribosome (grey). Adapted from PDB ID: 5LKS (Myasnikov et al., 2016) (B) Chemotherapy treatments blocks protein synthesis. Immunoblot analysis of puromycin incorporation into nascent proteins following treatment of K562 cells with 2.5µM AUR, 1µM 2ME, 8.3µM CPT, 7.1µM BLE, 6.0µM 5FU, 0.5mM SUL or cycloheximide (CHX, 1hr) for 12 hrs. (C) C22 in RPL18A and C39 in RPL37A are modified following treatment with multiple chemotherapies. (D) Summary of top-scoring genes and corresponding pathways in different locations that promote resistance or sensitivity to AUR treatment. (E) Cellular heatmap displaying the location of genes that mediate resistance and sensitivity to AUR treatment. (F) Inhibition of GSH biosynthesis sensitizes cells to AUR. Top, K562-dCas9-KRAB cells expressing the indicated sgRNAs were treated with 1 µM AUR or vehicle control. Bottom, K562 cells were co-treated with 1 µM AUR and 50 µM BSO. Relative proliferation was determined after 96 hrs. by measuring cellular ATP concentrations. ***p< 0.0001.

**Figure S4:**
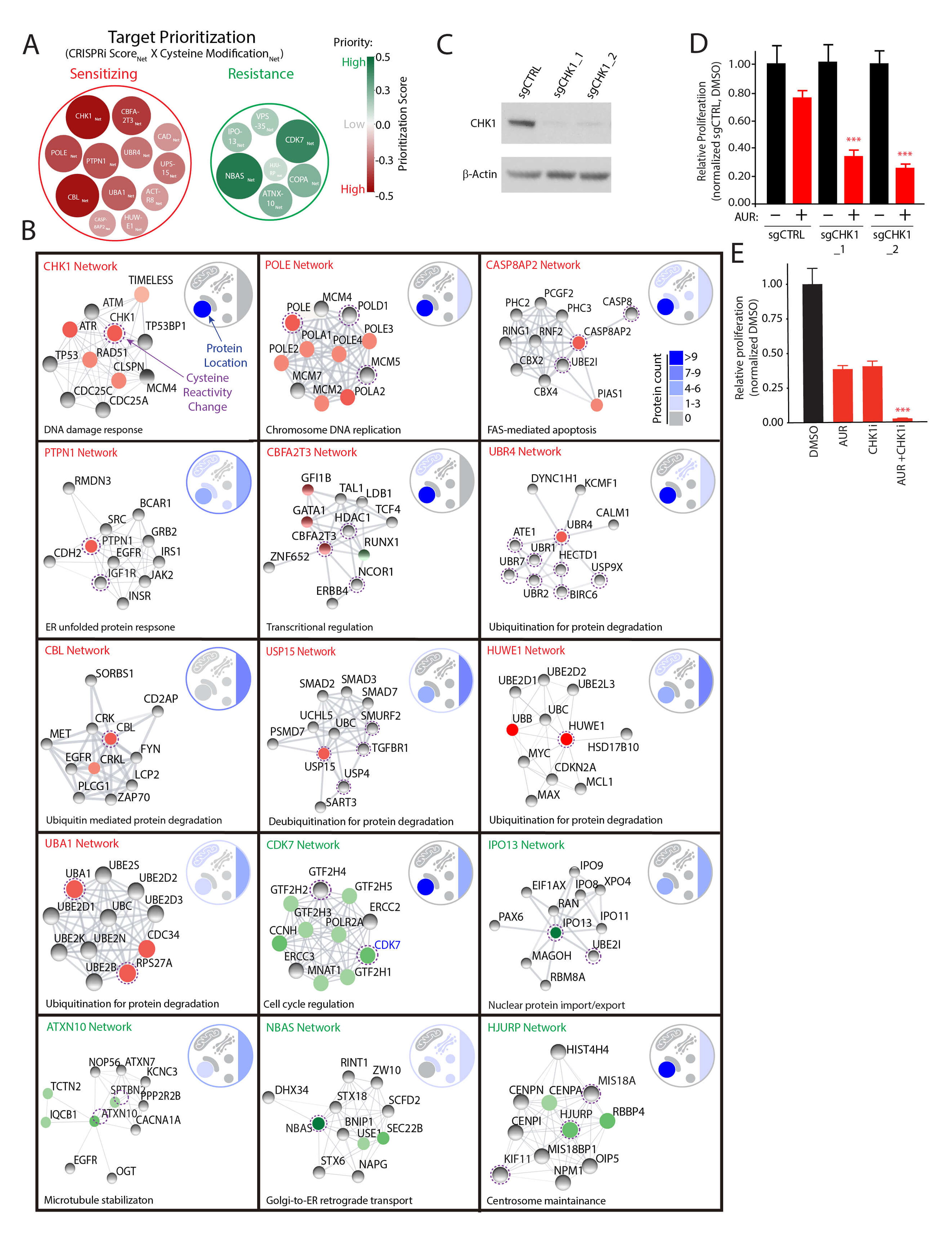
Proteogenomic prioritization of AUR targets. (A) Prioritization of targets based on the CRISPRi score and cysteine reactivity changes in their corresponding STRING interaction network (see also **Supplemental Table 4**). (B) Network analysis of functional targets for AUR and corresponding pathways and cellular locations. (C) Immunoblot analysis of CHK1 levels in K562-dCas9-KRAB cells expressing the indicated sgRNAs. (D) Depletion of CHK1 potentiates AUR-mediated proliferation. Relative proliferation of K562- dCas9-KRAB cells stably expressing the indicated sgRNAs following treatment with 1 µM AUR was determined by measuring cellular ATP concentrations following 96 hrs. of treatment. (E) CHK1 inhibition potentiates AUR cytotoxicity. Proliferation of K562 cells treated with 0.5µM CHK1i or 1µM AUR was measured following 96 hrs. as described in (D).

**Figure S5:**
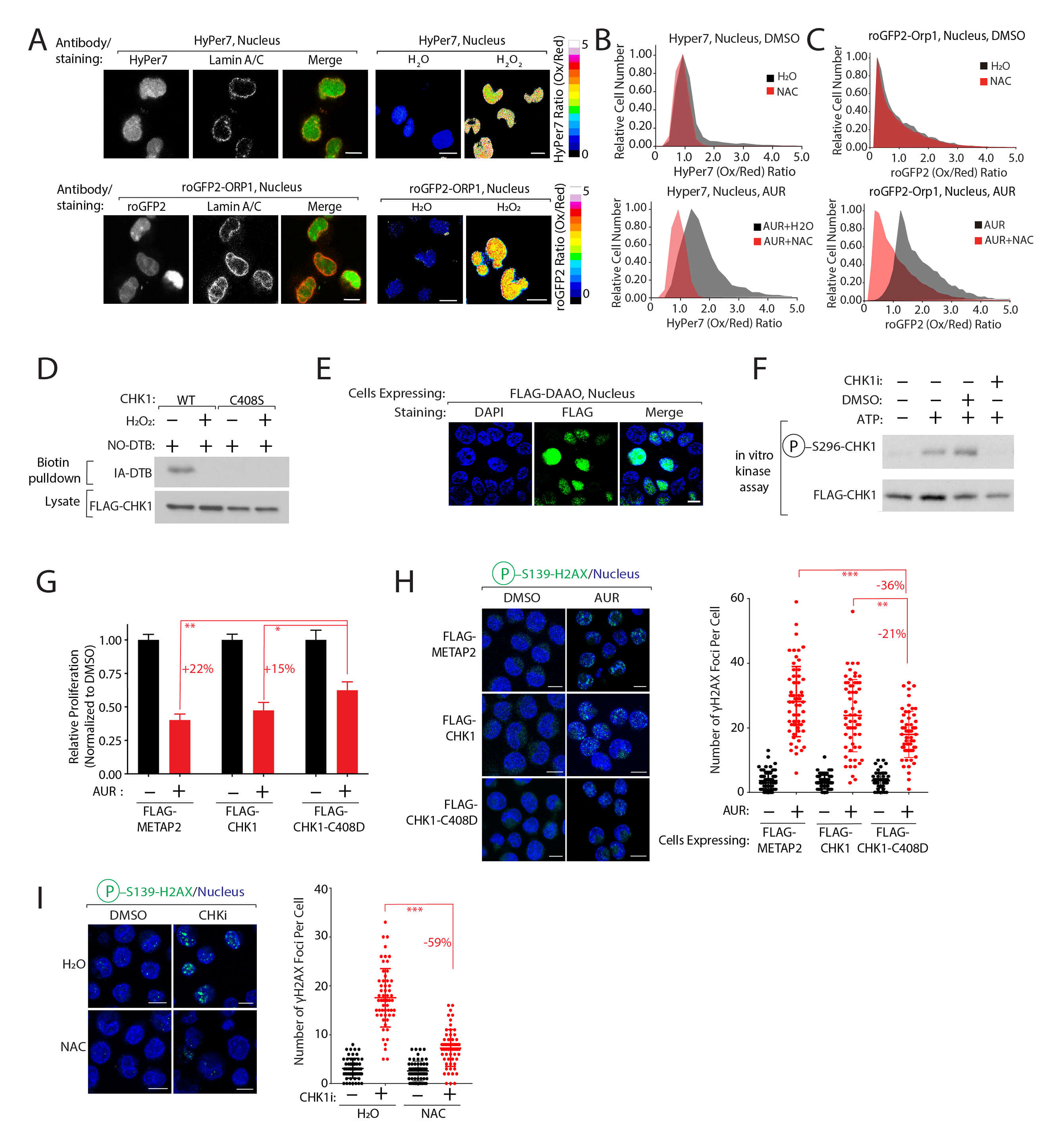
Cysteine 408 in CHK1 functions as a nuclear H_2_O_2_ sensor. (A) Left, Immunofluorescence analysis of K562 cells stably expressing the HyPer7 or roGFP2-ORP1 reporters in the nucleus, co-stained with Lamin A/C. Right, Immunofluorescence analysis of K562 cells stably expressing the HyPer7 or roGFP2-ORP1 reporters in the nucleus following H_2_O or 100 µM H_2_O_2_ treatment for 5 min. The ratiometric image was constructed by taking the ratio of the emission intensity of Oxidized (Ox) vs Reduced (Red) HyPer7 or roGFP2-ORP1. (B-C) Addition of NAC decreases nuclear H_2_O_2_ levels following AUR treatment. K562 cells expressing nuclear HyPer7 (B) or roGFP2-ORP1 (C) were treated with the indicated compounds for 6 hrs (5 mM NAC, 1.5 µM AUR) and the Ox/Red ratio of HyPer7/roGFP2-Orp1 was determined by flow cytometry (see also Figure 2C). (D) H_2_O_2_ treatment blocks alkylation of CHK1•C408. HEK-293T cells stably expressing CHK1 or CHK1•C408S were treated with 100 µM H_2_O_2_. IA-DTB modified CHK1 was determined by immunoblot following enrichment from cell lysates treated with a cysteine-specific IA-DTB. (E) Immunofluorescence analysis of FLAG-DAAO localized to nucleus in HEK-293T cells. (F) Characterization of CHK1 in vitro kinase assay. Immunoblot analysis of CHK1 in-vitro kinase assay following the addition of 5 µM CHK1 kinase inhibitor (CHK1i). (G) CHK1-C408D protects cells from AUR cytotoxicity. Relative proliferation of K562 cells stably expressing the indicated proteins was determined by measuring the changes in cellular ATP concentrations following treatment with 0.5 µM AUR for 96 hrs. (H) Left, immunofluorescence analysis of K562 cells stained with H2AX•S139 and DAPI. K562 cells stably expressing the indicated proteins were treated with 0.5 µM AUR for 24 hrs. Right, quantification of H2AX•S139 in the indicated samples. (I) NAC protects cells from CHK1i mediated DNA damage. Left, immunofluorescence analysis of K562 cells stained with H2AX•S139 and DAPI. K562 cells were co-treated with 2 µM CHK1i and 5 mM NAC for 24 hrs. Right, quantification of H2AX•S139 in the indicated samples. \**P* < 0.05, \*\**P* < 0.001, \*\*\**P* < 0.0001. Scale bar=10 µm.

**Figure S6:**
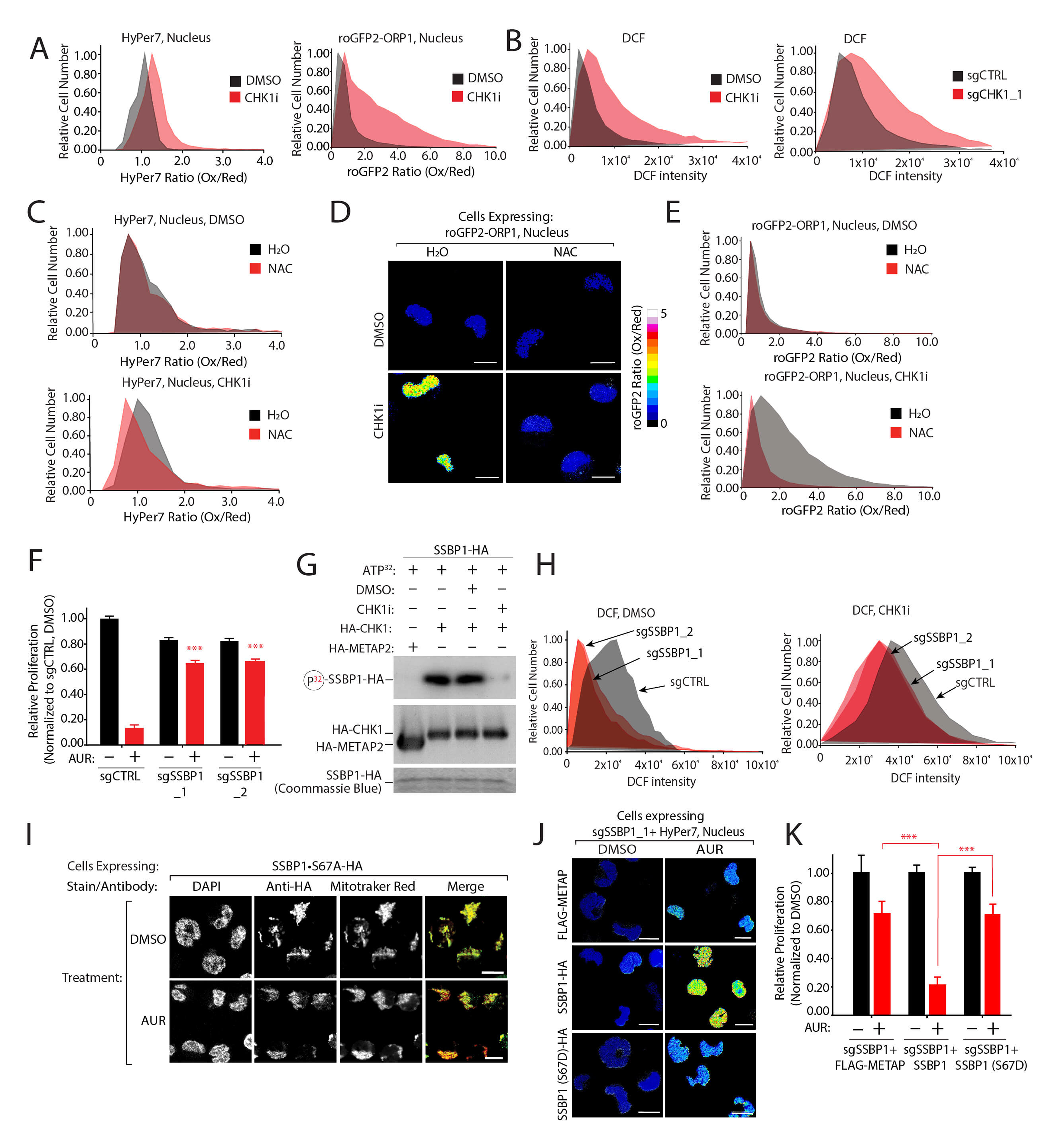
CHK1 regulates nuclear H_2_O_2_ levels through SSBP1 phosphorylation. (A) CHK1 inhibition increases nuclear H_2_O_2_. K562 cells expressing nuclear HyPer7 (left) or roGFP2-ORP1 (right) were treated for 48 hrs. with 2µM CHK1i. The Ox/Red ratio for each reporter was determined by flow cytometry. (B) Inactivation of CHK1 increases DCF fluorescence. Flow cytometry was used to monitor changes in DCF fluorescence in K562-dCas9-KRAB cells treated with CHK1i (2 µM) or expressing the indicated sgRNAs. (C) NAC treatment reverts CHK1- mediated nuclear H_2_O_2_ levels. K562 cells expressing nuclear localized HyPer7 were treated with 2µM CHK1i and 5mM NAC. Nuclear H_2_O_2_ levels were determined following 48 hrs of treatment as described in (A). (D) CHK1 inhibition increases nuclear H_2_O_2_ levels. K562 cells expressing nuclear roGFP2-ORP1 were treated with CHK1i (2 µM) and NAC (5 mM) for 48 hrs and the Ox/Red ratio of roGFP2-ORP1 was determined by immunofluorescence. (E) Flow cytometry analysis of nuclear localized roGFP2-ORP1 (Ox/Red) ratio following inhibition of CHK1. K562 cells expressing nuclear roGFP2-ORP1 were treated as in (D) and analyzed as in (A). (F) SSBP1 depletion rescues AUR cytotoxicity. K562 cells expressing the indicated sgRNAs were treated with 1.5 µM AUR or vehicle and proliferation was determined by measuring cellular ATP concentrations following 96 hrs of treatment. (G) CHK1i blocks the phosphorylation of SSBP1. CHK1 in vitro kinase assay was conducted with SSBP1-HA and purified HA-CHK1 or HA- METAP2 (control protein). CHK1 was treated with 5µM CHK1i and its phosphorylation of SSBP1 was monitored autoradiogram analysis. (H) SSBP1 functions downstream of CHK1 to regulate DCF fluorescence. K562-dCas9-KRAB cells expressing the indicated sgRNAs were treated with CHK1i or vehicle control and analyzed as described in (A). (I) The localization of SSBP1•S67A phosphodeficient mutant is not regulated by CHK1. Immunofluorescence analysis of K562 cells expressing SSBP1•S67A-HA following treatment with 1.5 µM AUR for 6 hrs. (J) SSBP1•S67D phosphomimetic mutant decreases nuclear H_2_O_2_. Reintroduction of SSBP1-HA or SSBP1•S67D- HA into K562 cells depleted of SSBP1. Cells were treated with 1.5 µM AUR or vehicle and nuclear H_2_O_2_ levels were measured with nuclear localized HyPer7 (see also Figure 3H). (K) The SSBP1•S67D mutant rescues AUR cytotoxicity. Cell lines described in (J) were treated with 1.5 µM AUR and proliferation was determined as described in (F). \*\*\**P* < 0.0001. Scale bar=10 µm.

**Figure S7:**
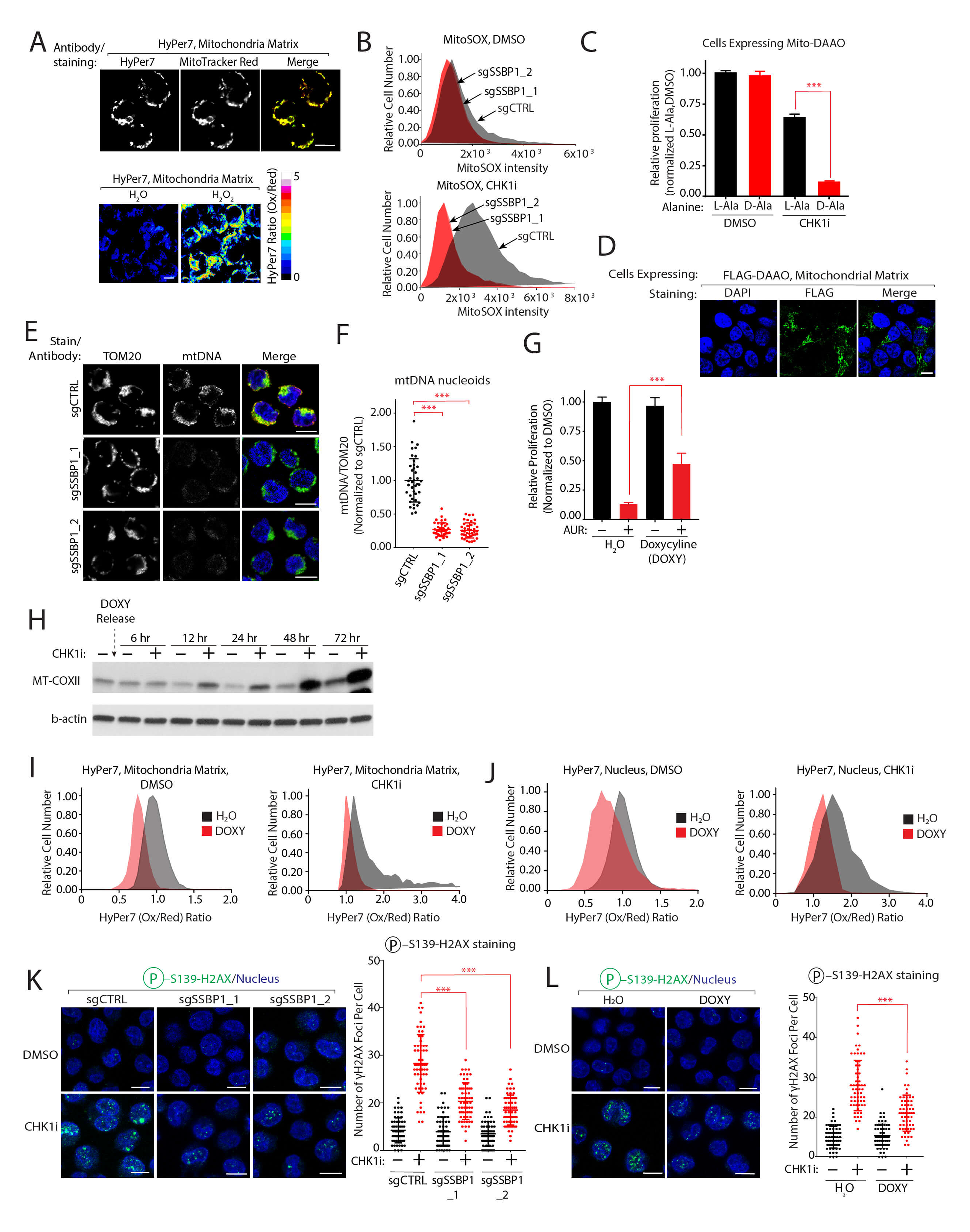
Mitochondrial translation regulates nuclear H_2_O_2_ levels. (A) Top, immunofluorescence analysis of K562 cells stably expressing the HyPer7 reporters in the mitochondrial matrix, co-stained with MitoTracker Red. Bottom, immunofluorescence analysis of K562 cells stably expressing the HyPer7 in the mitochondrial matrix following H_2_O or 100 µM H_2_O_2_ treatment for 5 min. The ratiometric image was constructed by taking the ratio of the emission intensity of oxidized HyPer7 over reduced HyPer7. (B) CHK1 regulates mitochondrial superoxide in a SSBP1-dependent manner. Mitochondrial superoxide was determined by monitoring changes in MitoSOX fluorescence with flow cytometry in K562-dCas9-KRAB cells expressing the indicated sgRNAs and treated with 2 µM of CHK1i for 48 hrs. (C-D) Mitochondrial H_2_O_2_ increases CHK1i cytotoxicity. K562 cells expressing mitochondrial matrix DAAO were treated with 1 µM CHK1 or 10 mM L-Ala or D-Ala, and proliferation was determined by measuring relative ATP concentrations following 96 hrs of treatment (C). Immunofluorescence analysis of FLAG-DAAO in K562 cells (D). (E-F) SSBP1 depletion decreases mtDNA content. Immunofluorescence analysis of mtDNA content in K562-dCas9-KRAB cells expressing the indicated sgRNAs (E). Quantification of mtDNA content following depletion of SSBP1 (F). (G) Doxycycline (DOXY) partially rescues AUR growth inhibition. K562 cells were co-treated with 2.3 µM DOXY and 1.5 µM AUR, and a relative change in proliferation was determined as described in (C). (H) CHK1 regulates mitochondrially encoded proteins. K562 cells were treated with 2.3 µM DOXY for 48 hrs, at which point the media was exchanged and replaced with 2 µM CHK1i or vehicle control. MT-COXII protein levels were determined by immunoblot (see also Figure 4F). (I) CHK1 functions upstream of mitochondrial translation to regulate mitochondrial H_2_O_2_ levels. K562 cells expressing HyPer7 localized to the mitochondrial matrix were treated with 2.3 µM DOXY for 24 hrs followed by 2 µM CHK1i for 48 hrs. The HyPer7 Ox/Red ratio was determined by flow cytometry. (J) CHK1 functions upstream of mitochondrial translation to regulate nuclear H_2_O_2_ levels. K562 cells expressing nucleus localized HyPer7 were treated with 2.3 µM DOXY for 24 hrs followed by 2 µM CHK1i for 48 hrs. Nuclear H_2_O_2_ levels were determined as in (I). (K-L) Genetic or pharmacological inhibition of mitochondrial translation blunts CHK1i mediated DNA-damage. Immunofluorescence analysis of H2AX•S139 staining in K562-dCas9-KRAB cells expressing the indicated sgRNAs targeting SSBP1 (K) or treated with 2.3µM DOXY (L) were treated with 2 µM CHK1i for 48 hrs (n= 60 cells counted per condition). \*\*\**P* < 0.0001. Scale bar=10 µm.

**Figure S8:**
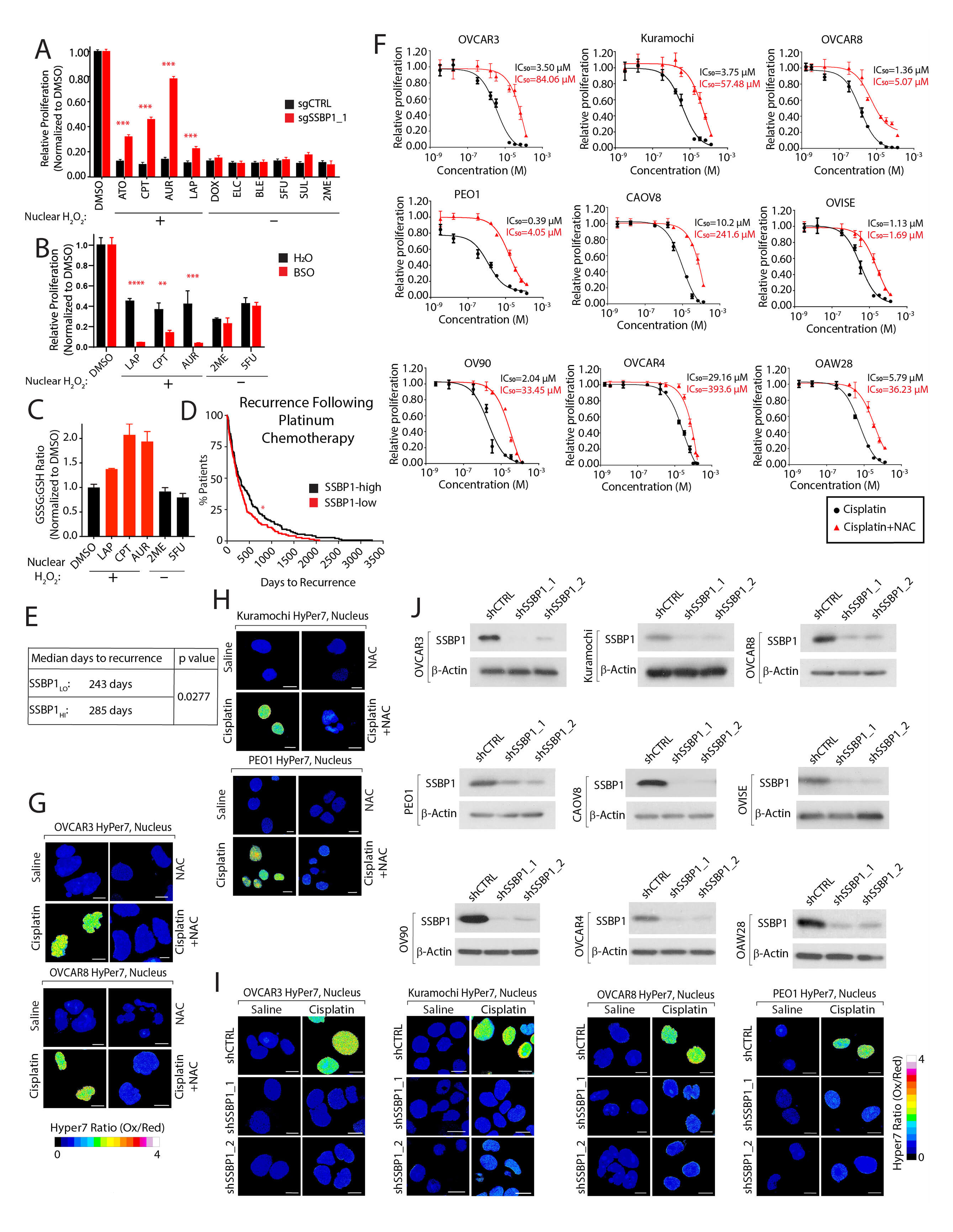
SSBP1 levels regulate platinum-based nuclear H_2_O_2_ and resistance in ovarian cancers. (A) SSBP1 depletion partially rescues the proliferation deficit due to chemotherapies that increase nuclear H_2_O_2_ levels. K562-dCas9-KRAB cells expressing the indicated sgRNAs were treated with each compound, and relative proliferation was determined by measuring cellular ATP concentrations following 96 hrs of treatment. (B) GSH depletion sensitizes cells to chemotherapy treatment that increases nuclear H_2_O_2_. K562 cells were co-treated with 50 µM BSO and the indicated compounds (1 µM LAP, 2.5 µM CPT, 0.25 µM AUR, 0.5 µM 2ME, 1 µM 5FU). Proliferation was determined as in (A). (C) The GSSG:GSH ratio increases following treatment with chemotherapies that increase nuclear H_2_O_2_. K562 cells were treated with the indicated compounds (1.5µM LAP, 8.3µM CPT, 2.5µM AUR, 1µM 2ME or 6.0µM 5FU) for 24 hrs, and the GSSG:GSH ratio was determined. (D-E) Patients with lower SSBP1 expression have shorter duration to tumor recurrence following platinum-based chemotherapy. High-grade serous ovarian patients were stratified based on SSBP1 mRNA expression. Clinical data were obtained from (Cancer Genome Atlas Research, 2011; Villalobos et al., 2018). (F) NAC treatment decreases cisplatin cytotoxicity in ovarian cancer cell lines. Cell lines were co-treated with 5mM NAC or vehicle control and the indicated concentrations of cisplatin. Relative proliferation was determined as described in (A). (G-H) Cisplatin increases steady state levels of nuclear H_2_O_2_ in ovarian cancer cells, which can be rescued by NAC treatment. Cell lines expressing nuclear HyPer7 were co-treated with 5mM NAC or vehicle control and 8.3 µM cisplatin for 24 hrs. The ratiometric image was constructed by taking the ratio of the emission intensity of oxidized HyPer7 over reduced HyPer7. (I) Immunoblot analysis of SSBP1 in ovarian cancer cell lines expressing the indicated shRNAs. (J) Depletion of SSBP1 in ovarian cancer cell lines decreases cisplatin regulated nuclear H_2_O_2_ levels. Cell lines co-expressing nuclear HyPer7 and the indicated shRNAs were treated with cisplatin and analyzed as described in (G-H). \**P* < 0.05, \*\**P* < 0.001, \*\*\**P* < 0.0001. Scale bar=10 µm.

